# Broad epigenetic shifts in the aging *Drosophila* retina contribute to its altered rhythmic transcriptome

**DOI:** 10.1101/2025.06.17.660160

**Authors:** Sarah E. McGovern, Gaoya Meng, Makayla N. Marlin, Sophia A. Pruitt, Vikki M. Weake

## Abstract

Alterations in biological rhythms are a common feature of aging, and disruption of circadian rhythms can exacerbate age-associated pathologies. The retina is critical for detecting light for both vision and for transmitting time-of-day information to the brain, synchronizing rhythms throughout the body. Disruption of circadian rhythms by manipulating the molecular clock leads to premature retinal degeneration in flies and mice, and gene expression rhythms are disrupted in models of age-associated ocular disease. Despite this, it is unknown how or why the gene expression rhythms of the retina change with age. Here, we show that ∼70% of the *Drosophila* transcriptome is rhythmically expressed throughout the diurnal cycle, with ∼40% of genes showing altered rhythms with age. These transcriptome-wide changes in aging photoreceptors are accompanied by shifts in the rhythmic patterns of RNA Polymerase II (Pol II) occupancy, histone H3 lysine 4 (H3K4) methylation, and chromatin accessibility, without major changes in occupancy of the circadian clock transcription factors Clock (Clk) and Cycle (Cyc). Instead, aging decreases genome-wide levels of several different histone methyl marks including H3K4 methylation, whose relative levels across the day correlate with the phase of rhythmic gene expression. Moreover, individual knockdown of the three H3K4 methyltransferases in young photoreceptors results in massive disruptions to rhythmic gene expression that resemble those observed during aging. We conclude that there are broad epigenetic shifts in the aging retina, including decreased histone methylation, that contribute to changes in biological rhythms even in the presence of a robust molecular circadian clock.

## INTRODUCTION

Biological rhythms are highly synchronized with the diurnal cycle, and are largely controlled by an environmentally-entrained molecular circadian clock that enables organisms to anticipate and respond to predictable day-night cycles through rhythmic gene expression programs (Patke et al. 2019). During aging, there are changes in circadian behavior and gene expression that are also observed in age-related pathologies (Hood & Amir 2017; Acosta-Rodríguez et al. 2021). Disrupted circadian rhythms have been linked to many human health conditions, including neurodegenerative diseases, cancer, and psychological disorders (Acosta-Rodríguez et al. 2021; Rijo-Ferreira & Takahashi 2019). In the retina, which senses light for vision and to entrain circadian rhythms (Sanes & Zipursky 2010), many critical biological processes such as neurotransmitter release and visual sensitivity are synchronized with the diurnal cycle (Tosini et al. 2008). Maintaining these proper biological rhythms is critical for health of the eye because loss of the circadian clock transcription factor *Bmal1* results in cataracts and other age-associated phenotypes in mice (Kondratov et al. 2006), while retina-specific *Bmal1* loss leads to accelerated age-associated photoreceptor degeneration (Baba et al. 2018). In addition, rhythmic gene expression is highly altered in mouse models of the age-associated ocular disease, diabetic retinopathy (Ye et al. 2023; Vancura et al. 2021). The circadian clock is highly conserved across species (Stanton et al. 2022), and in flies, photoreceptor-specific expression of the dominant negative circadian clock transcription factor Clk results in retinal degeneration (Jauregui-Lozano et al. 2022; Hodge et al. 2022). Although the retina transcriptome is highly rhythmic in young mice (Storch et al. 2007), and both mouse and *Drosophila* retinal cells undergo vast age-dependent transcriptomic changes (Marola et al. 2025; Hall et al. 2017), it is unclear whether aging alters the overall levels of gene expression or simply changes its rhythm i.e., alters the timing of gene expression with respect to the diurnal cycle. The latter possibility is supported by observations that many genes exhibit age-dependent changes in rhythmic gene expression in both flies and mice (Solanas et al. 2017; Kuintzle et al. 2017).

The photoreceptors in the retina are the most abundant molecular clock-regulated cells in the *Drosophila* head and are peripherally involved in entraining circadian behaviors to light (Yoshii et al. 2015), although central clock cells in the brain can also be entrained directly by Cryptochrome and specialized Rhodopsins (Schlichting 2020). *Drosophila* photoreceptors bear several functional similarities to the intrinsically photosensitive retinal ganglion cells (ipRGCs) involved in photoentrainment in mammals, which utilize an invertebrate-style phototransduction cascade initiated by the Rhodopsin-like melanopsin (Graham et al. 2008; Hattar et al. 2003). In flies, the molecular clock is composed of the transcription factors, Clk and Cyc, that activate the expression of rhythmic output genes including ones encoding their own repressors, Period (Per) and Timeless (Tim), generating an approximately 24-hour rhythmic pattern of gene expression (Patke et al. 2019). In addition, light itself can regulate gene expression independent of the molecular clock because many genes in *Drosophila*, particularly those involved in phototransduction, remain rhythmically expressed in the *tim* mutant (Wijnen et al. 2006). We previously found changes in chromatin accessibility around the binding motifs for Clk and Cyc in old photoreceptors in flies (Jauregui-Lozano et al. 2022), suggesting that circadian-regulated gene expression might change with age. However, the nature of these potential rhythmic gene expression changes and the molecular mechanisms involved were not identified.

Epigenetic alterations, including changes to chromatin accessibility and histone modifications, are a hallmark of aging that contribute to age-associated changes in gene expression (López-Otín et al. 2023). The transcription factor network established by the circadian clock coordinates with epigenetic marks including histone modifications to establish rhythmic gene expression (Sahar & Sassone-Corsi 2013; Koike et al. 2012). There are daily rhythmic patterns of Pol II occupancy (Xu & Li 2023), chromatin accessibility (Yuan et al. 2024), and levels of active and repressive histone modifications (Zhu & Belden 2020). Although rhythmic, the timing of deposition of these epigenetic patterns does not correlate with the peak of rhythmic gene expression genome-wide (Le Martelot et al. 2012; Trott & Menet 2018; Koike et al. 2012). In the aging retina, there are epigenetic alterations to DNA methylation (Corso-Díaz et al. 2020), chromatin accessibility (Xu et al. 2022; Jauregui-Lozano et al. 2023), and histone methylation (Jauregui-Lozano et al. 2023), suggesting that epigenetic changes could impact the rhythmic transcriptome.

Here, we used *Drosophila* retinal photoreceptors as a model to explore the rhythmic gene expression and chromatin changes during aging, focusing on this single light-sensing cell type that undergoes light-dependent degeneration upon disruption of biological rhythms (Jauregui-Lozano et al. 2022). We show that many genes undergo age-related changes in their daily rhythmic expression, which are largely independent of any changes in their overall expression levels across the day. We identify broad changes in the rhythmic diurnal occupancy of Pol II, chromatin accessibility, and histone methylation during aging, and identify genome-wide decreases in levels of multiple histone methyl marks in old photoreceptors. Last, we show that knockdown of H3K4 methyltransferases disrupts rhythmic gene expression in young photoreceptors, overlapping with many of the rhythm changes observed during aging. Our data demonstrate that the molecular clock is insufficient to explain the rhythmic changes during aging, and suggest that instead epigenetic shifts in the aging retina contribute to altered biological rhythms and dampened oscillations under the diurnal cycle.

## RESULTS

### Aging reshapes the photoreceptor rhythmic transcriptome

To understand how photoreceptor aging impacts rhythmic gene expression across the diurnal cycle, we performed photoreceptor nuclear RNA-seq across a standard day-night cycle in young (D10) and old (D50) flies (Fig. 1a). Principal component analysis (PCA) separated samples both by the time of day, zeitgeber time (ZT), and by age, with decreased ZT separation in old flies (Fig. 1b). We identified age-dependent changes in rhythmic gene expression using dryR, a package that employs a statistical framework to detect differential rhythmicity between multiple conditions (Weger et al. 2021). When comparing rhythmic gene expression between young and old flies, 8180 genes met the statistical cutoff for accurate model fitting. Of these categorized genes, 74% are rhythmic in young and/or old flies: 28% have the same rhythm at both ages, 19% lose rhythmicity, 10% gain, and 13% undergo changes in rhythmic parameters such as phase or amplitude (Fig. 1c,d; Table S1). Only 26% of expressed genes are arrhythmic at both ages, indicating that the majority of photoreceptor genes are rhythmically expressed in a normal daily light cycle. Restricting this analysis to intronic reads instead of exonic reads, we obtained highly similar results, suggesting these age-dependent changes in rhythmic gene expression represent alterations in pre-mRNA levels rather than steady-state mRNA, likely reflecting transcription (Fig. S1).

**Figure 1:**
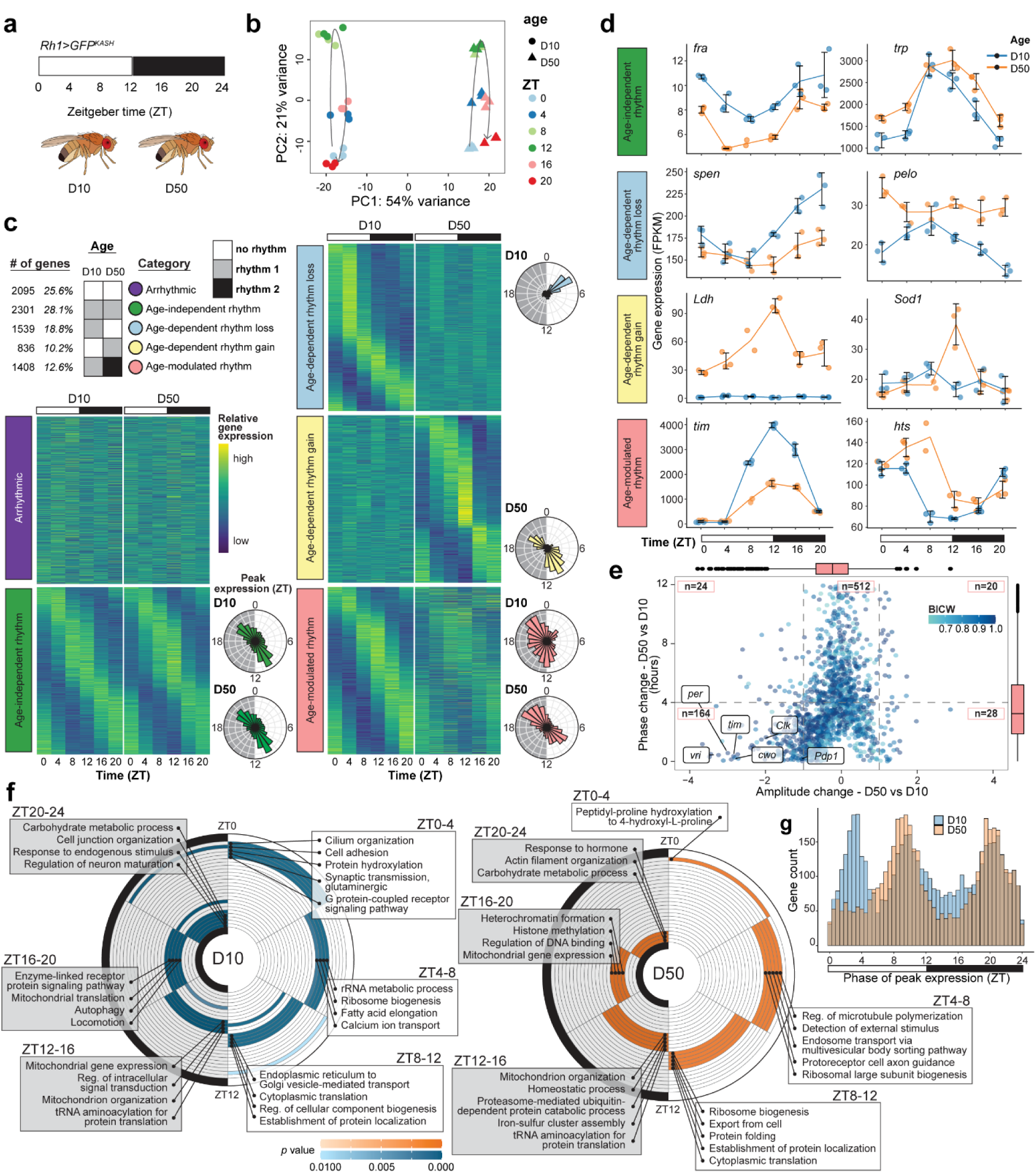
Aging alters the rhythmic photoreceptor transcriptome. **a,** Diurnal photoreceptor nuclear RNA-seq analysis in young (D10) and old (D50) *Rh1>GFP^KASH^* flies (*n* = 3). **b,** PCA of RNA-seq samples based on variance-stabilized gene expression levels. **c,** Rhythmic dryR exon-based categories (BICW ≥ 0.6). Heatmaps show *z* score of relative expression, and radial histograms show phase of peak gene expression. **d,** Line plots of representative genes. Individual FPKM data points shown with line representing mean ± s.d. **e,** Scatterplot showing age-dependent amplitude versus phase change, colored by BICW with selected core clock genes labeled. Box-and-whisker plots represent the IQR (box) and median (line) values. **f,** Representative GO terms for rhythmic genes expressed most highly across four-hour intervals at D10 versus D50. Terms are colored by *p* value. **g**, Histogram showing phase of rhythmic gene expression at D10 and D50.

We previously found that expression of dominant negative Clk in photoreceptors resulted in significant retinal degeneration that could be rescued by raising the flies in constant darkness (Jauregui-Lozano et al. 2022). Similarly, *period* mutants that disrupt the clock exhibit light-dependent retinal degeneration (Fig. S2a,b). Despite the role of the core clock in maintaining photoreceptor health, the rhythmicity of core clock gene expression is not entirely disrupted in old flies, as observed by others (Luo et al. 2012; Rakshit et al. 2012). Instead, the core clock genes predominantly fall into the age-modulated rhythm category and exhibit a steep drop in expression amplitude without a concurrent change in phase, indicating their expression timing remains in phase with the day-night cycle (Fig. 1d-e). In addition, clock genes exhibit an overall decrease in gene expression in old flies, which is not a typical feature for other genes in the age-modulated rhythm category that shows increased expression and phase shifts (Fig. S3a-b; Fig. 1e). Interestingly, unlike *per* mutants, *Clk, Cyc,* and *tim* mutants did not show significant retinal degeneration by D20 (Fig. S2a), suggesting that not all disruptions to the molecular clock are detrimental to photoreceptors.

In general, rhythmic changes in expression are not associated with a coincident change in overall gene expression levels (Fig. S3a). An exception is the genes that gain rhythmicity with age, 15% of which also increase in overall expression levels. These genes include many of the previously-described “late-life cyclers” (Kuintzle et al. 2017), whose rhythmic expression is induced by oxidative stress. Though we do see oxidative stress response genes including *superoxide dismutase 1* (*Sod1*) and *lactate dehydrogenase* (*Ldh*) gain rhythmicity in old photoreceptors (Fig. 1d), a larger proportion of these age-dependent rhythm gain genes are involved in visual perception, determination of adult lifespan, and cytoplasmic translation (Fig. S3c). Over the course of the diurnal cycle, photoreceptor cells carry out an elegantly orchestrated gene expression program (Fig. 1f, S3d). In young flies, genes involved in neuronal structure and function including synaptic transmission shows a peak expression at the cusp of the light phase in preparation for light exposure and perception. Midday, as the photoreceptors continue to sense light, there is a peak in expression of genes with functions in calcium channel activity, fatty acid metabolism, and ribosome biogenesis. At the end of the day and into the early night, photoreceptors express genes involved in mitochondrial function, translation, protein folding, and protein localization establishment, along with genes encoding structural components of the rhabdomere. During the late night, photoreceptors express autophagy and carbohydrate metabolism genes, before once again preparing for the onset of light exposure.

Intriguingly, genes with different phases of peak expression in young flies change differently during aging. Many genes that lose rhythmicity during aging peak in expression between ZT0-4 in young flies (Fig. 1c,g). In young flies, there are three times of day when rhythmic genes most often reach their peak expression: ZT0-4, 8-12, and 20-24 (Fig. 1g). In old flies, this trimodal distribution converts to a bimodal one since the ZT0-4 peak is lost. Notably, the genes that peak at this time in young flies include 69 transcription factors, 52 of which lose this peak in expression in old flies, suggesting stark shifts in transcriptional regulation during aging (Table S1). These genes also include factors involved in cell adhesion and cellular structure, which may contribute to the increase in retinal degeneration we observe in aging flies (Escobedo et al. 2023). Rhythmic expression plots for any genes of interest, similar to those presented in Fig. 1d, can be visualized at: https://vikkiweake.shinyapps.io/shinyr_vweake/. Together, these data demonstrate that aging results in widespread changes to rhythmic transcription that impact more than a third of expressed genes, including loss, gain and modulation of rhythms.

### Aging alters the rhythms of Pol II occupancy, chromatin accessibility, and H3K4 methylation

To determine how chromatin structure might contribute to altered rhythmic transcription, we examined the patterns of Pol II occupancy, chromatin accessibility, and H3K4 methylation during aging. To do this, we profiled H3K4me1, H3K4me2, and H3K4me3 individually, together with Pol II every four hours in young (D10) and old (D50) photoreceptors using CUT&RUN. We also examined chromatin accessibility in these same samples using ATAC-seq. We focused initially on H3K4 methylation because H3K4me3 contributes to Pol II initiation and promoter-proximal release (Shilatifard & Org 2012; Wang et al. 2023), and the TRX/MLL1 methyltransferase that deposits this modification has a role in clock-regulated gene activation at certain genes (Zhang et al. 2022; Katada & Sassone-Corsi 2010). We observed a distinct diurnal pattern of Pol II occupancy, H3K4 methylation, and chromatin accessibility at individual genes including *split ends* (*spen*, Fig. 2a) and genome-wide (Fig. 2b). In young flies, Pol II occupancy reaches a maximum genome-wide at promoters at ZT8 and ZT20, four hours prior to the light turning on or off. Between these times, at ZT12, chromatin accessibility reaches a maximum at promoters. In old flies, Pol II still has two waves of maximal occupancy, but the timing is shifted four hours earlier to ZT4 and ZT16. Maximum chromatin accessibility similarly shifts back four hours in old flies, but also spreads across multiple time points in contrast to its temporally-restricted peak of accessibility at ZT12 at D10 (Fig. 2b).

**Figure 2:**
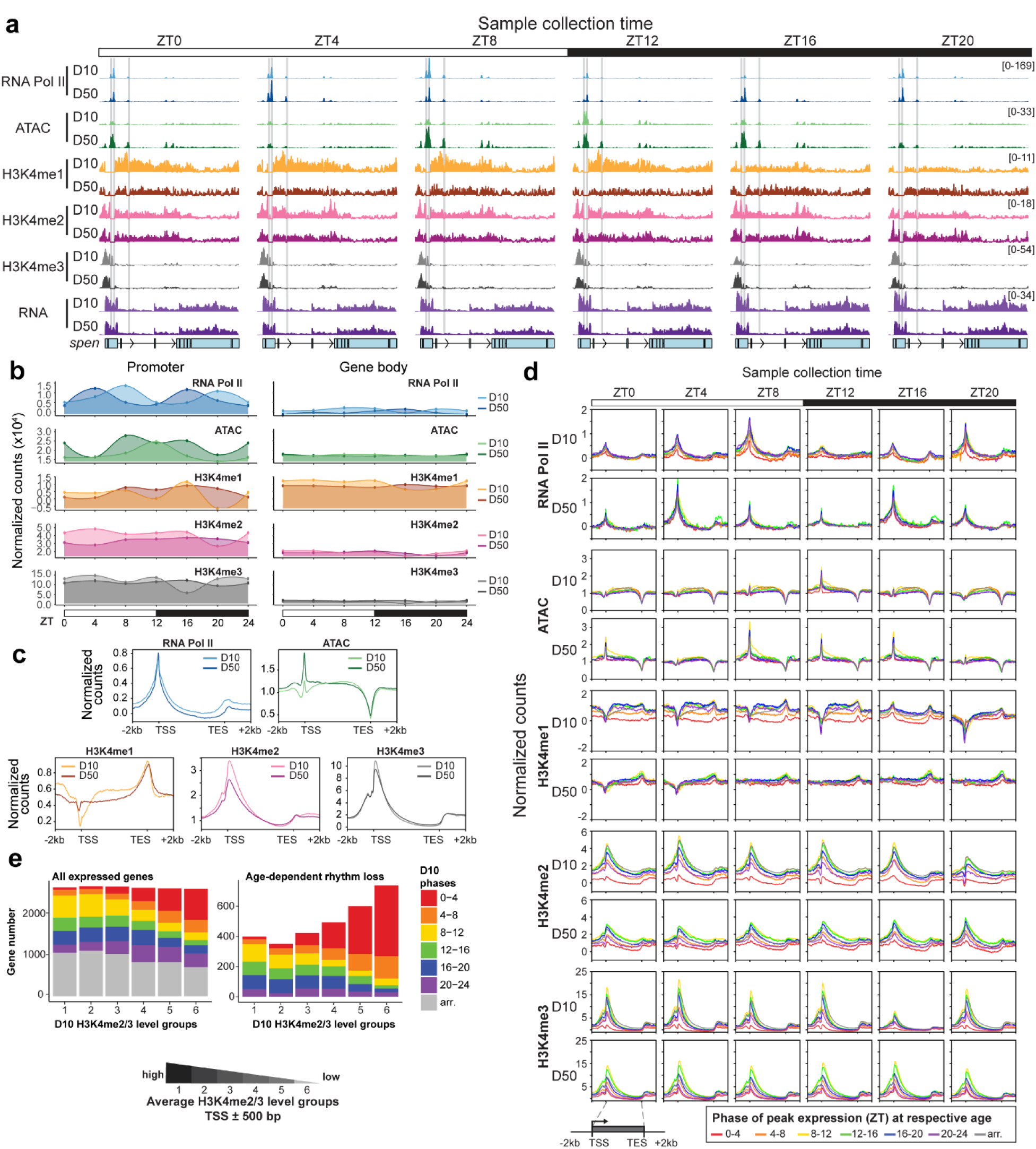
Aging alters rhythmic Pol II, chromatin accessibility, and H3K4 methylation. **a,** ATAC-seq, RNA-seq, and IgG-normalized CUT&RUN tracks at *split ends* (*spen*). Data are mean (*n* = 3). Identified Clk and Cyc peaks are shown as horizontal grey lines. **b,** Line plots of averaged signal at promoters (transcription start site, TSS ± 500 bp) or gene bodies (500 bp downstream of TSS to transcription end site, TES) of all expressed genes. **c,** Gene metaplots representing mean normalized counts across all ZTs for all expressed genes at D10 versus D50. **d,** Gene metaplots representing mean normalized counts separated by phase of peak expression at each ZT at D10 or D50. Genes are clustered based on the phase of peak expression at D10 or D50 determined by nuclear RNA levels at each age (ZT: 0-4, 4-8, 8-12, 12-16, 16-20, 20-24, arr. = arrhythmic). **e,** Bar plot (left) showing the phase distribution of all expressed genes based on their relative H3K4me2 and H3K4me3 levels at D10. Genes were separated into six H3K4me2/3 level groups ranging from highest (group 1) to lowest (group 6) based on their average normalized H3K4me2/3 scores at TSS ± 500 bp. Bar plot (right) showing only genes that lose rhythmicity during aging, based on their phase distribution at D10.

H3K4 methylation also shows dynamic patterns in young flies that are lost during aging (Fig. 2a,b). Unlike the periodic oscillation of Pol II occupancy and chromatin accessibility, promoter H3K4me3 levels remain high across most of the day in young flies, dropping only once during the night at ZT16, accompanied by a transient increase in H3K4me1. Four hours later, at ZT20, promoter H3K4me3 levels rise accompanied by decreased H3K4me1/2 preceding the start of the light phase. H3K4me1 and H3K4me2 are also present at gene bodies for most of the day, dipping only once at ZT16. Overall, in young flies H3K4 methylation levels remain stably high during the day and decrease only briefly at night. In contrast, in old flies, these changes in all three H3K4 methyl marks during the night are ablated (Fig. 2b). In addition, there is an overall decrease of promoter H3K4me2/3 levels accompanied by an increase of H3K4me1 at transcription start sites at D50 (Fig. 2c). We previously observed similar decreases in H3K4me2/3 in old photoreceptors at a single time point by ChIP-seq, controlling for nucleosome occupancy using bulk histone H3 (Jauregui-Lozano et al. 2023). Thus, the decrease in H3K4me2/3 methylation detected by CUT&RUN does not reflect decreased nucleosome occupancy. Moreover, this age-dependent decrease in H3K4me2/3 is unlikely caused by deficient Pol II recruitment because the overall promoter Pol II signal increases slightly in old flies, though there is a moderate decrease on the gene body, potentially indicative of deficient elongation (Fig. 2c). Overall, aging alters the dynamic patterns of Pol II occupancy, chromatin accessibility, and H3K4 methylation genome-wide in photoreceptors.

### H3K4 methylation patterns correspond to the phase of rhythmic transcription

Surprisingly, we observed a near complete disconnect between the genome-wide rhythms of Pol II occupancy, chromatin accessibility, and the timing of rhythmic gene expression. For example, the gene *spen* shows peak expression at ZT18 in young flies (Fig. 1d), whereas its maximum Pol II occupancy occurs at ZT8, and chromatin accessibility peaks at ZT12 (Fig. 2a,b). Clustering genes by the time of their peak rhythmic expression shows that the overarching diurnal pattern of Pol II and chromatin accessibility is the same across nearly all genes irrespective of peak expression time (Fig. 2d, Fig. S4a). Even arrhythmic genes show highest levels of Pol II occupancy at ZT8 and 20, and chromatin accessibility at ZT12 in young flies (Fig. 2d, grey lines). At most genes, H3K4 methylation also exhibits the same pattern, regardless of when a gene peaks in expression. One exception is the small number of core circadian clock genes including *per*, *Clk*, and *vri*, that show highly rhythmic patterns of H3K4me3, corresponding to their peak expression, suggesting that these genes may have a unique mode of regulation (Fig. S5). However, at the vast majority of photoreceptor genes, the diurnal patterns of Pol II occupancy, chromatin accessibility, and H3K4 methylation are independent of rhythmic expression timing at either age. Since our nuclear RNA-seq primarily reflects pre-mRNA levels, the observed discrepancy between transcript oscillation and epigenetic rhythms is unlikely to result from post-transcriptional regulatory steps that affect mRNA abundance.

Instead, we observe a marked difference in the relative signal levels for H3K4 methylation between rhythmic genes that peaked at different times of the day (Fig. 2d, Fig. S4a). In both young and old flies, rhythmic genes that peaked between ZT8-16, spanning the transition from day to night, had much higher H3K4me2/3 levels relative to other groups of rhythmic genes. Conversely, genes that peaked in expression at the beginning of the day, from ZT0-4, had much lower levels of H3K4me2/3. For H3K4me1, genes peaking between ZT8-20 had the highest levels, with much lower levels for genes that peaked between ZT0-4. In old flies, this separation of H3K4me2/3 levels is retained, though slightly reduced in magnitude, but H3K4me1 levels are no longer distinguishable between groups. These data suggest that the phase of expression for many rhythmically transcribed genes may be partially dictated by their relative level of H3K4 methylation. If so, we would expect that relative H3K4 methylation levels could predict the phase of rhythmic gene expression. When we separated genes into six groups based on their relative levels of H3K4me2/3, we find that genes with the lowest levels of H3K4me2/3 (group 6) are more often expressed at ZT0-4, and also more often lose rhythmicity during aging (Fig. 2e). Conversely, rhythmic genes with the highest H3K4me2/3 levels (groups 1 - 3) are more often expressed between ZT8-16, and tend to gain or maintain their rhythmicity during aging, especially when compared with group 6 (Fig. S4b,c). Although H3K4 methylation often correlates with gene expression levels, these differences in H3K4me2/3 levels between the genes that peak at different times of day do not simply reflect lower expression (Fig. S4d). Instead, we wondered if differences in H3K4 methylation levels might be reflective of broader changes in the chromatin state, dictated by transcription factor networks. The basic helix-loop-helix (bHLH) transcription factors Clk and Cyc form a heterodimer to activate the expression of circadian-regulated genes (Patke et al. 2019), and we previously showed that disrupting Clk activity alters the expression of 20% of photoreceptor genes and changes global chromatin accessibility (Jauregui-Lozano et al. 2022). We therefore asked whether the age-dependent changes in Pol II occupancy, chromatin accessibility, and H3K4 methylation result from changes in Clk:Cyc binding.

### The molecular clock does not explain age-dependent changes in rhythmic transcription

To identify changes in Clk:Cyc binding in aging photoreceptors, we profiled GFP-tagged Clk and Cyc in young and old photoreceptors using CUT&RUN at two time points (ZT12 and ZT16) flanking their peak of DNA binding, relative to the trough, ZT4 (Abruzzi et al. 2011; Rivas et al. 2021) (Fig. 3a). We identified 281 Clk:Cyc peaks based on the presence of either Clk or Cyc at any age or ZT (Fig. 3b; Table S2). As a control, we confirmed that there was no detectable signal at these regions in GFP CUT&RUN from untagged flies (*w^1118^*). As expected, there is higher enrichment of Clk:Cyc at ZT12 and ZT16 relative to ZT4 in both young and old flies, consistent with the maintained phase of core clock gene expression observed by RNA-seq. Clk and Cyc are both enriched at all peaks, indicating there are no sites uniquely targeted by either transcription factor, consistent with Clk and Cyc binding as a heterodimer (Darlington et al. 1998). Additionally, there are no new Clk:Cyc peaks in old flies, suggesting that Clk:Cyc do not gain any new targets during aging. Though no new peaks are observed in old flies, average Clk signal at its peaks decreases at D50 despite Cyc remaining similar at both ages (Fig. 3c). This phenomenon is more apparent for peaks with lower levels of Clk to begin with at D10, while peaks with high levels of Clk retain strong signal at D50 and have increased Cyc signal. Many of these strongest Clk:Cyc targets (Fig. S6a,b, cluster 1) correspond to core clock genes. Since Clk:Cyc bind DNA as a heterodimer, our data do not support a loss of Clk:Cyc occupancy at most target genes in old flies. Instead, we interpret these differences in Clk:Cyc CUT&RUN signal to reflect changes in accessibility of the epitope at different ages and/or times, potentially due to association with other proteins.

**Figure 3:**
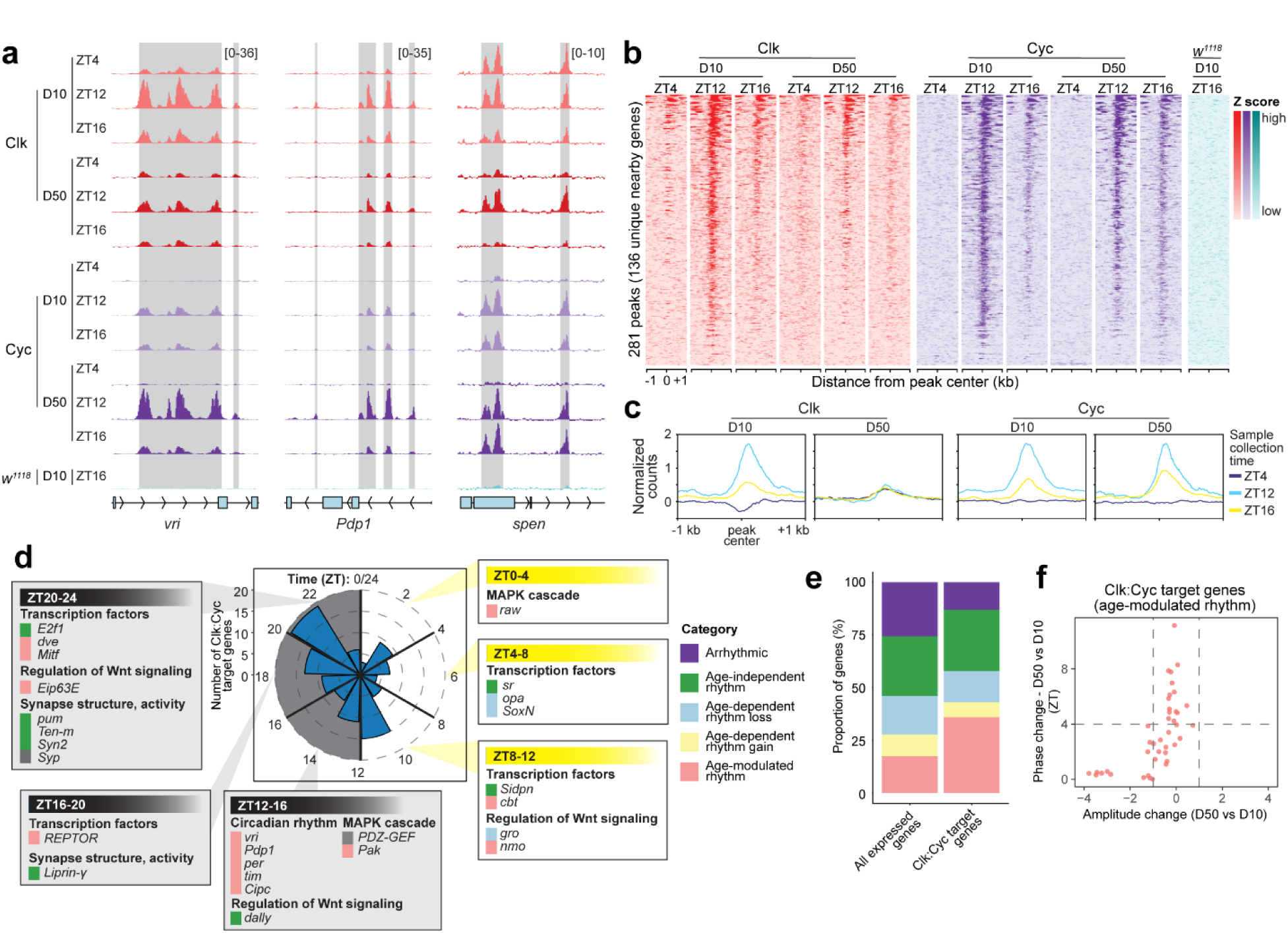
Clk and Cyc occupancy does not correlate with age-dependent rhythmic changes. **a**, IgG-normalized CUT&RUN tracks of Clk and Cyc for indicated genes at each age and ZT with peaks represented by grey boxes. Data are mean (*n* = 3). **b,** Heatmaps representing all Clk and Cyc peak centers ± 1 kb colored by relative signal (*z* score) across the same region (rows). **c,** Gene metaplots representing mean normalized counts for Clk:Cyc peaks at each ZT and age. **d,** Radial histogram showing peak gene expression time for Clk:Cyc target genes at D10 with representative GO terms illustrated for rhythmic genes expressed most highly across selected four-hour intervals. Boxes adjacent to gene names are colored by age-dependent rhythmicity category. **e,** Bar plot of proportion of Clk:Cyc target genes compared with all expressed genes, colored by age-dependent rhythmicity category. **f,** Scatterplot showing age-dependent amplitude versus phase change, for Clk:Cyc target genes in the age-modulated rhythm category.

The Clk:Cyc peaks map to 136 nearby genes, similar in number to recent ChIP-seq analysis of Clk in heads (Rivas et al. 2021), but an order of magnitude lower than previous studies (Abruzzi et al. 2011; Meireles-Filho et al. 2014). These Clk:Cyc target genes include the well-characterized core clock genes, as well as many other genes encoding transcription factors, Wnt signaling regulators, components of the MAPK cascade, and factors important for neuronal function (Fig. 3d). Clk:Cyc target genes are more often rhythmic as compared with all expressed genes (Fig. 3e; Fig. S6c), and have higher chromatin accessibility, Pol II enrichment, and more dramatic oscillations in H3K4 methylation levels throughout the day (Fig. S6d). Though some Clk:Cyc target genes are expressed in the same phase (around ZT12) as the core clock genes like *per* and *tim*, others are expressed at different times across the circadian day, especially later into the night and sometimes even in the early morning (Fig. 3d). This disparity in timing of Clk:Cyc occupancy and target gene expression timing suggests that additional factors are necessary for the ability of Clk and Cyc to stimulate transcription at specific target genes. We note that some Clk:Cyc peaks exhibit Clk signal even at ZT4, consistent with the idea that, at least in photoreceptors, Clk:Cyc activity may not be completely restricted to ZT12-16 (Fig. S6b). Although Clk:Cyc targets tend to be rhythmic, they undergo a similar variety of changes in rhythmicity during aging when compared with all expressed genes (Fig. 3e). For instance, while the core clock genes bound by Clk:Cyc (*per*, *tim*, *vri*) all show decreased amplitude with age, many other Clk:Cyc-bound genes that are not components of the core molecular clock change in phase, or even lose, or gain rhythmicity during aging (Fig. 3e-f). Thus, the decrease in Clk signal at many Clk:Cyc peaks at D50 does not correspond to any one type of rhythmicity change, and Clk:Cyc target genes behave similarly to the general trend we observe for all genes. Based on this, our data indicate that changes in the recruitment of Clk:Cyc do not explain the majority of the rhythmic gene expression changes that occur during aging. Instead, because Clk:Cyc target genes exhibit age-dependent decreases in H3K4me2/3 levels similarly to all other genes, we propose that the ability of Clk:Cyc to stimulate transcription requires additional co-activators that modulate chromatin state. This suggests that changes that impact the chromatin state in old flies should impact Clk:Cyc activity in the same way as any other transcription factor.

### H3K4 methyltransferases are required for rhythmic gene expression

Because H3K4 methylation levels correlate with the phase of gene expression for rhythmic genes, we hypothesized that the age-dependent decrease in H3K4 methylation in photoreceptors contributes to the sweeping changes in rhythmic transcription we observe in old flies. In *Drosophila*, there are three H3K4 methyltransferases: SET domain containing 1 (Set1), Trithorax (Trx), and Trithorax-related (Trr) (Tie et al. 2014). All three methyltransferases are necessary for H3K4me3 because ubiquitous expression of RNAi against Set1 in larvae decreases bulk H3K4me3 levels relative to non-specific mCherry RNAi control (Fig. 4a, Fig. S7a), while CUT&RUN from photoreceptors expressing RNAi against Trx or Trr shows that both contribute to H3K4me3 at promoters (Fig. 4b). In addition, Trx knockdown, but not Trr, decreases H3K4me2 levels at promoters and H3K4me1 levels on the gene body. To identify genes that required each of these H3K4 methyltransferases for rhythmic expression, we profiled the diurnal photoreceptor transcriptome of D10 flies expressing RNAi against Set1, Trx, and Trr, relative to mCherry RNAi control (Fig. 4c). Using dryR, we obtained 52 potential mathematical models for this four-condition comparison representing genes with altered rhythmicity in at least one genotype (Table S3). Six of these models contained at least 200 genes above statistical criteria (see methods; Fig. 4d). Supporting a critical role for these H3K4 methyltransferases in rhythmic gene expression, only 263 genes (2%) retained the same rhythm upon knockdown of any of the H3K4 methyltransferases (Fig. 4d, unchanged rhythm). In contrast, over 3033 genes (26%) lost rhythmicity in at least one of the H3K4 methyltransferase knockdowns, while more than 1000 genes exhibit altered rhythmic parameters (Fig. 4d, Fig. S7b). Notably, over two thirds of the genes that lost rhythmicity were affected by Trx or Trr knockdown, but not Set1, indicating a shared but non-redundant role for Trx and Trr in regulating rhythmic expression of many target genes (Fig. 4d-f). Conversely, 500 genes gained rhythmicity uniquely in the Set1 knockdown (Fig. 4d; Table S3). Similar to our observations for the aging data, differential rhythmic expression does not correspond to overall differential gene expression (Fig. S7c), with the exception of genes in the Trx-and Trr-modulated rhythm categories that exhibit a tendency towards decreased overall expression. Thus, in *Drosophila,* all three H3K4 methyltransferases are required non-redundantly for H3K4me3 and rhythmic gene expression across most of the transcriptome. Since disruption of circadian rhythms either in the *period* mutant or upon photoreceptor-specific expression of dominant negative Clk results in retinal degeneration (Fig. S2; (Jauregui-Lozano et al. 2022)), the dysregulated rhythmic gene expression caused by H3K4 methyltransferase knockdown might also increase susceptibility to retinal degeneration. Indeed, we previously showed in an RNAi screen that photoreceptor-specific knockdown of either Set1 or Trr results in premature retinal degeneration (Escobedo et al. 2023), and here we also observed significant premature retinal degeneration in Set1 knockdown flies compared with mCherry control at D30 (Fig. 4g). Together, these data indicate that the age-dependent decrease in H3K4me2/3 levels (Fig. 2c) may have a profound impact on the rhythmic transcriptome.

**Figure 4:**
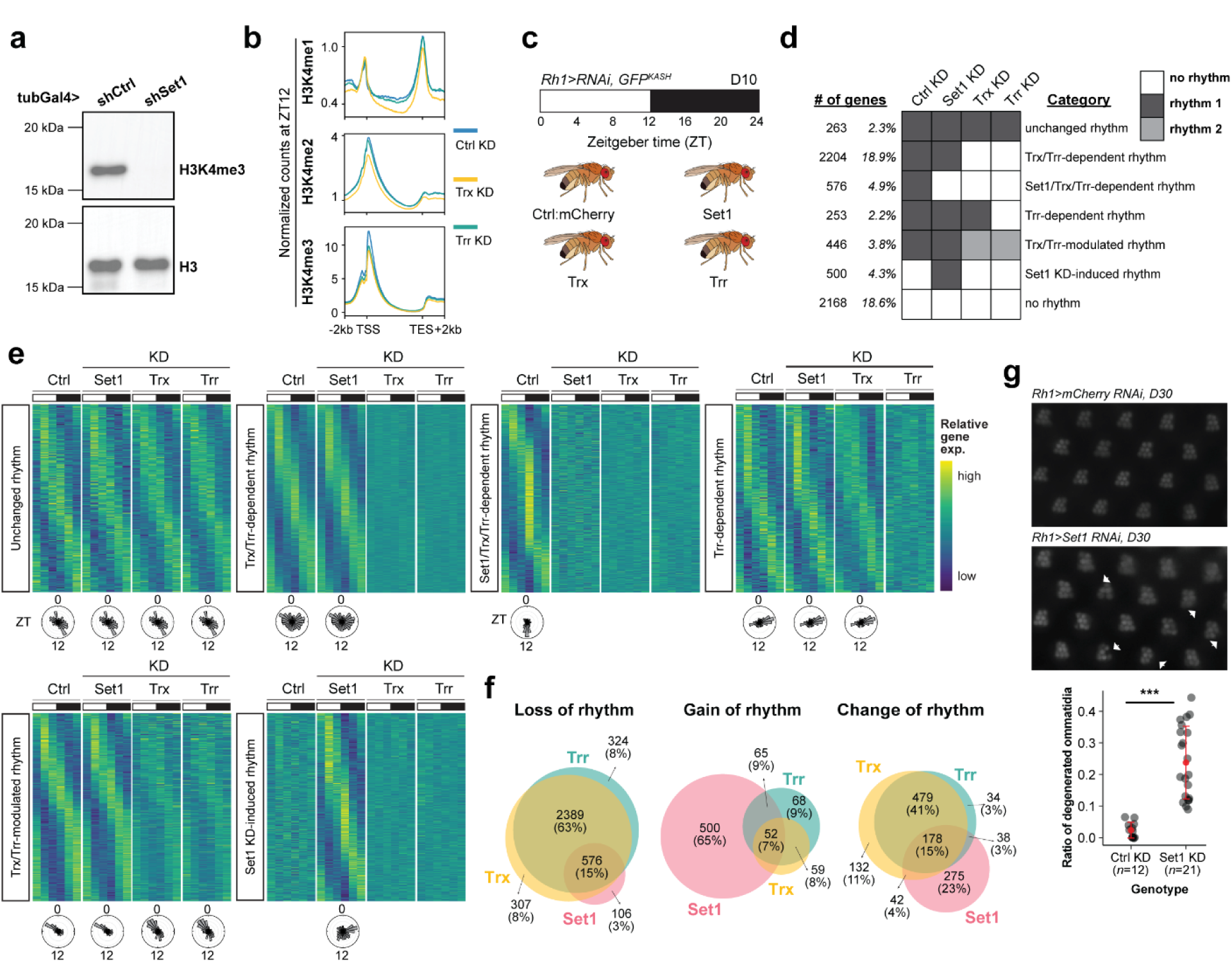
Knockdown of H3K4 methyltransferases disrupts rhythmic expression. **a,** Western blot of H3K4me3 in larvae expressing *tub>RNAi* against control (mCherry RNAi) or Set1. Histone H3 is shown as a loading control on a separate blot. **b,** Gene metaplots representing mean normalized counts for CUT&RUN of each histone methyl mark at ZT12 for all expressed genes from photoreceptors expressing *Rh1>RNAi* against control, Trx, or Trr (*n* = 3). **c,** Diurnal photoreceptor nuclear RNA-seq analysis in young (D10) flies (*n* = 3) expressing *Rh1>RNAi* against control (mCherry), Set1, Trx, or Trr. **d,** Number of genes categorized into rhythmicity groups based on differential rhythmicity across all four conditions (BICW ≥ 0.4) for categories with >200 genes. Heatmaps for each category show *z* score of relative expression, and radial histograms show phase of peak gene expression. **e,** Venn diagrams showing the number of genes overlapping for the loss, gain, and change of rhythm categories for each H3K4 methyltransferase knockdown. **f,** Retinal degeneration was assessed by optical neutralization in male D30 flies expressing *Rh1>RNAi* against control (mCherry RNAi) or Set1. Flies were raised in 12-hr:12-hr light:dark (L:D). Number of biological replicates (*n*) is indicated per condition. *** *p* < 0.001, Student’s t-test.

### Knockdown of H3K4 methyltransferases recapitulates many of the age-dependent changes in rhythmic gene expression

To determine if decreased H3K4me2/3 methylation in aging photoreceptors could at least partially account for the sweeping changes in rhythmic gene expression in old photoreceptors, we compared the rhythmicity changes observed in the H3K4 methyltransferase knockdowns with those in aging photoreceptors (Table S3). Since the two experiments used different control lines, we restricted our analysis to only genes that were similar in both controls (i.e., arrhythmic in both mCherry RNAi and D10 or rhythmic in both) (Fig. S8a). 80% of the genes that lose rhythmicity during aging also lose rhythmicity in at least one of the H3K4 methyltransferase knockdown lines, mostly due to Trx or Trr (Fig. 5a,b, Fig. S8b). Similarly, more than 40% of the genes with age-modulated rhythms also show modulated rhythms upon knockdown of the H3K4 methyltransferases. Decreased amplitude is a common feature of genes that overlap between the aging- and knockdown-modulated categories (Fig. 5c), suggesting that H3K4 methylation is important for maintaining the amplitude of rhythmic gene expression. In fact, many of the core molecular clock components such as *per, Cipc,* and *cwo* exhibit amplitude decreases in both aging and upon knockdown of all three H3K4 methyltransferases (Fig. 5d). Notably, 59% of the age-modulated genes exhibit total loss of rhythmicity in at least one of the H3K4 methyltransferase knockdowns (Fig. S8b), suggesting that in some cases, decreased amplitude resulting from diminished H3K4 methylation could entirely abolish rhythmicity. We note that a few genes show discordant rhythmicity changes between the individual H3K4 methyltransferase knockdowns e.g., loss in Trr and Trx versus modulation in Set1, leading to differences in the total numbers of genes in some comparisons (Fig. S8b).

**Figure 5:**
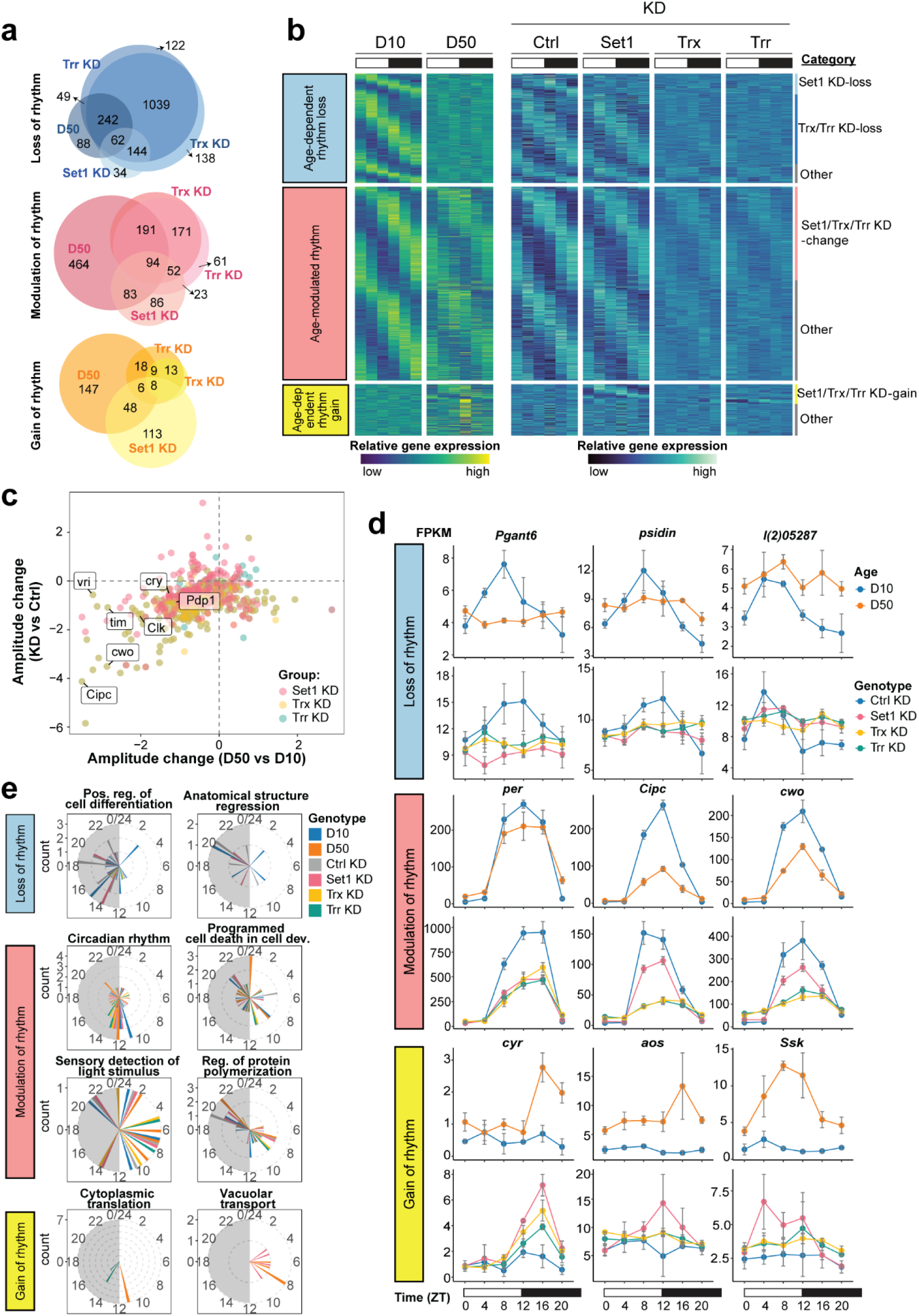
Knockdown of H3K4 methyltransferases resembles aging. **a,** Venn diagrams showing the number of genes overlapping for the loss, gain, and change of rhythm categories for each H3K4 methyltransferase knockdown versus aging. **b,** Heatmaps showing *z* score of relative expression for genes (rows) at D10 and D50, and in the knockdown control, Set1, Trx, or Trr RNAi samples at D10. All genes in the age-dependent loss, modulated, and gain categories are shown, clustered within each category by rhythmicity change in each knockdown (vertical lines to the right of the heatmap). **c,** Scatterplot comparing change in rhythmic gene expression amplitude during aging versus histone methyltransferase knockdown. Select core clock genes are labeled. **d,** Line plots of gene expression levels for representative genes in the overlapping categories. Data are mean FPKM ± s.d. (*n* = 3). **e,** Selected GO term analysis of genes in each overlapping aging/knockdown rhythmicity category displayed as radial histograms indicating peak gene expression phase in each sample.

Interestingly, several genes involved in autophagy completely lose rhythm in both aging and histone H3K4 methyltransferase knockdown (Fig. 5e), such as *Autophagy-related 9 (Atg9), Autophagy-related 18a (Atg18a), Death-associated APAF1-related killer (Dark),* and *Death regulator Nedd2-like caspase (Dronc)*. In addition, genes involved in the response to light exhibit altered rhythmic expression in both aging and upon methyltransferase knockdown. Strikingly, a portion of genes whose rhythmicity are induced with age also gain rhythmicity upon Set1 knockdown (Fig. 5a,b). Apart from “late-life cyclers” previously described as being involved in the oxidative stress response (Kuintzle et al. 2017), these “gain-of-rhythm” genes encode proteins involved in translation, largely components of the ribosome (Fig. 5e).

Together, these data indicate that the genome-wide decreases in H3K4me2/3 levels in aging flies contribute to age-dependent changes in rhythmic transcription – even at core circadian clock genes. To test if alleviating the age-dependent decrease of H3K4me2/3 levels would restore amplitude of these core clock genes, we expressed RNAi against the H3K4me3 demethylase Kdm5 (Secombe et al. 2007) in photoreceptors and examined transcript levels by qRT-PCR at D50. Although photoreceptors constitute the most abundant clock cell in the head, we did not observe a significant difference in *per, tim,* or *vri* transcript levels at the peak of their expression (ZT12) between Kdm5 knockdown and control (Fig. S8c), despite the efficiency of knockdown in this Kdm5 RNAi line (Fig. S8d). Thus, long-term knockdown of the H3K4 demethylase Kdm5 is not sufficient to rescue the age-dependent changes in rhythmic gene expression, indicating that there might be broader epigenetic changes in old flies that act together to alter rhythmicity – not only H3K4 methylation.

### Multiple histone methyl marks decrease genome-wide in aging photoreceptors

We previously found that old photoreceptors also have genome-wide decreases in H3K36me3 (Jauregui-Lozano et al. 2023), hinting at more widespread changes in the aging photoreceptor epigenome. To examine this possibility, we used CUT&RUN to profile H3K4me2/3 and H3K36me3, and two repressive marks, H3K9me3 and H3K27me3 in young (D10) and late middle-aged (D40) photoreceptors (Fig. 6a). Similar to our previous findings, H3K36me3 decreases over gene bodies in older flies, together with decreased H3K4me2/3 at promoters, albeit to a lesser extent than by D50. These alterations in histone methylation are not restricted to active marks because both H3K9me3 and H3K27me3 also decrease with age. The decrease in histone methylation does not result from photoreceptor loss because these flies show no retinal degeneration at D40 (Hall et al. 2017), and we obtain similar yields for both RNA-seq and CUT&RUN samples at both young and old ages (data not shown). We also do not observe overall trends in gene expression *i.e.*, decreases in expression of H3K4 methyltransferases or increased expression of demethylases, that provide a simple explanation for the observed age-dependent decreases in histone methylation (Fig. S8e); for instance, both Set1 and Trx are actually upregulated in old photoreceptors at the transcript level.

**Figure 6:**
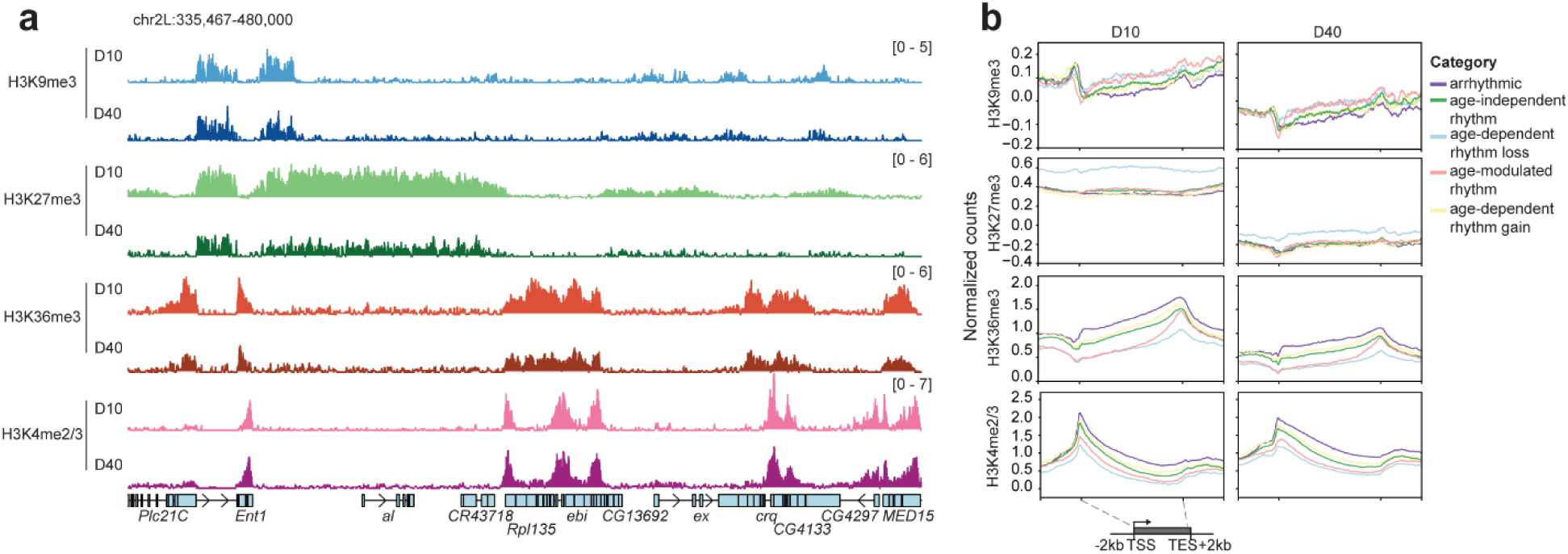
Histone methyl marks decrease genome-wide in aging photoreceptors. **a,** IgG-normalized CUT&RUN tracks of histone methyl marks for indicated genomic region at each age at ZT12. Data are mean (*n* = 4). **b,** Gene metaplots for age-dependent rhythmic categories.

Despite the unanswered question as to why histone methylation decreases with age, our data support a model in which these broad genome-wide changes in histone methylation during aging have combinatorial impacts on rhythmic transcription. When we examine the relative levels of each histone methyl mark at genes that show different changes in rhythmicity with age, we find that those genes that lose (*blue line*) or change (*red line*) rhythmicity during aging are marked by lower levels of H3K4me2/3 and H3K36me3 relative to the other categories of genes that are arhythmic, or that gain or maintain rhythmicity with age (Fig. 6b, S8f). In addition, the genes that lose rhythmicity with age also exhibit higher levels of H3K9me3 and H3K27me3 relative to other categories at both D10 and to a lesser extent at D40. These data indicate that the genes that lose rhythmicity with age may lie near repressive chromatin domains, marked by lower H3K4 methylation and higher levels of H3K37me3. Intriguingly, these same genes that lose rhythmicity with age and have the lowest levels of H3K4 methylation tend to be expressed in the early morning between ZT0-4 (Fig. 2e). Our data suggest that the broader chromatin state, defined in part here by combinatorial histone methylation levels, contributes both to the phase of rhythmic gene expression in young flies *<u>and</u>* to how aging will alter rhythmicity. Thus, the widespread epigenetic changes in old photoreceptors may play a major role in reshaping its rhythmic transcriptome.

## DISCUSSION

Here, we show that under normal daily light conditions, the rhythmic photoreceptor transcriptome is completely reshaped during aging at both the transcriptional and epigenomic level. Although the molecular clock has an important role in regulating rhythmic gene expression, and is a major regulator of the photoreceptor transcriptome (Jauregui-Lozano et al. 2022), our data indicate that changes in the synchronization of the molecular clock are insufficient as a causative explanation for the age-dependent alterations in rhythmic transcription. First, the phase of core clock genes remains synchronized with the light-dark cycle in old flies, indicating that photoentrainment remains intact in the old retina. Second, Clk and Cyc bound genes exhibit a similar age-dependent rhythmicity changes as compared to all other genes. Third, while genetic manipulations that strongly disrupt circadian rhythms (Clk dominant negative and the *period* mutant) result in light-dependent retinal degeneration, null mutations in several other core clock genes do not. Importantly, our analyses were performed in the context of standard light-dark cycles, and light itself can induce rhythmic gene expression patterns independent of the core clock machinery (Yoshii et al. 2015). These data suggest that age-dependent changes in biological rhythms in the retina are associated with widespread restructuring of the epigenome, even in the presence of a robustly synchronized core clock.

How might the epigenetic changes in aging photoreceptors reshape its rhythmic transcriptome? We, like others, observe decreases in the amplitude of core molecular clock genes such as *per* and *tim* during aging (Luo et al. 2012; Rakshit et al. 2012), which seems to be a hallmark of aging across circadian-regulated tissues. We propose that this decrease in the amplitude of core clock expression in aging photoreceptors results from an impairment in the kinetics of transcription activation due to reduced H3K4 methyltransferase activity. Supporting this idea, knockdown of the H3K4 methyltransferases in young flies decreases amplitude of core clock genes similarly to aging, and H3K4me2/3 levels are decreased genome-wide in old photoreceptors. We speculate that the overlap we observe between H3K4 methyltransferase knockdown and aging may be due to the importance of H3K4 methylation for productive transcription elongation, which in turn is critical for rhythmic transcription. In mouse embryonic stem cells, acute depletion of H3K4me3 decreases release of promoter-proximally paused Pol II, leading to decreased transcription elongation (Wang et al. 2023), while in plants, removal of H3K4me1 decreases Pol II processivity (Menon et al. 2024). Previous studies have attributed the observed disconnect between Pol II recruitment and the phase of gene expression to delays in post-transcriptional processing (Le Martelot et al. 2012). However, in mouse liver, levels of Pol II on the gene body, but not at promoters, correlates with transcription of clock-regulated genes (Zhu et al. 2018). Our data reveal that the relative levels of H3K4 methylation, rather than oscillating temporal patterns, correspond to the peak of transcription with the dynamic patterns of Pol II, chromatin accessibility, and H3K4 methylation occurring genome-wide, regardless of when a gene is maximally expressed. One possibility is that higher H3K4me2/3 levels enhance the rate of Pol II release and promote transcription elongation leading to a peak of expression between ZT8 – 12, while genes with lower H3K4me2/3 levels have decreased Pol II elongation and delayed expression. This model remains to be tested since we have not analyzed transcription elongation directly in aging photoreceptors due to the technical challenges associated with isolating photoreceptor nuclei from flies. Moreover, although we show that H3K4 methylation is a major contributor to maintaining rhythmic transcription in old flies, both non-histone substrates of the methyltransferases, and the other histone methyl marks that decrease in old flies could also impact circadian rhythms. Butterfly TRX has been shown to influence circadian rhythms by promoting the interaction between Clk and Per through arginine methylation of heat shock protein 68 (Zhang et al. 2022). Further, H3K9me3 and H3K27me3 methyltransferases impact circadian gene expression in mammals (Duong & Weitz 2014; Liu et al. 2024), and the H3K36 methyltransferase Set2 contributes to age-dependent splicing in photoreceptors (Jauregui-Lozano et al. 2023). Thus, decreases in methylation of multiple histone and non-histone substrates during aging could affect both transcription itself, and downstream processes such as splicing and RNA processing, leading to combinatorial and widespread impacts on rhythmic gene expression.

Our study leaves open the question of what causes the widespread changes in chromatin, particularly the decrease in histone methylation, in the aging retina. We do not observe expression changes in methyltransferases or demethylases at the transcript level that provide a simple explanation for the decrease in histone methylation, suggesting that another mechanism is likely involved. One potential explanation is that there are age-dependent changes in primary metabolism, which generate many of the substrates and cofactors required for histone modifications (Serefidou et al. 2019). Supporting this hypothesis, our prior metabolomic analysis identified substantial changes in metabolism in the aging *Drosophila* eye (Hall et al. 2021). Further, in other tissues such as mouse liver, dietary interventions that alter metabolism such as caloric restriction, which promotes lifespan, also increase the number of rhythmically expressed genes (Sato et al. 2017). Thus, the changes in metabolism that occur as a consequence of aging could provide one explanation for why circadian rhythms become dysregulated in old age. Since tissues receive distinct metabolic inputs that shape the epigenome, aging could impact rhythmic gene expression in distinct ways depending on the response of each tissue’s epigenome.

## METHODS

### Fly stocks and genetics

*Drosophila* were maintained on standard fly food at 25°C under 12-h:12-h light:dark conditions where ZT0 indicates the beginning of the light phase. Male flies were collected at indicated age (days post-eclosion, D) ± 1 day and ZT ± 15 minutes with dark phase collections (ZT12-20) performed under red light. *Rh1-Gal4>UAS-GFP^KASH^* flies were collected for aging RNA-seq and CUT&RUN/ATAC-seq studies (D10, D50). CUT&RUN studies of histone methyl marks were also performed in *Rh1-Gal4>UAS-GFP^KASH^* (D10, D40). For knockdown RNA-seq and CUT&RUN studies (D10), *Rh1-Gal4>UAS-GFP^KASH^* were crossed to either control (mCherry) RNAi, Set1 RNAi, Trx RNAi, or Trr RNAi. Clk and Cyc CUT&RUN (D10, D50) was performed in progeny from *Rh1-Gal4>UAS-GFP^KASH^* crossed to either GFP-tagged Clk or Cyc (Kudron et al. 2018), or *w^1118^*(untagged) control. A full list of fly stocks used in this study is provided in Table S4.

### Assessment of retinal degeneration

Retinal degeneration was assessed in at least 10 individual live male flies by bright-field microscopy using optic neutralization (Franceschini & Kirschfeld 1971). Images were de-identified and scored blindly for rhabdomere loss.

### qRT-PCR analysis of gene expression

Relative expression of indicated genes was examined using qRT-PCR in pupae in triplicate as previously described (Stegeman et al. 2018). cDNA was generated from 300 ng of RNA, and expression values were determined by qPCR using a reference curve relative to reference gene (*RpL32* and *eIF1A*). Primers are described in Table S4.

### Nuclei immuno-enrichment (NIE)-based RNA-seq, CUT&RUN, or ATAC-seq

Photoreceptor nuclei were immuno-enriched from *Rh1-Gal4>UAS-GFP^KASH^*heads using GFP antibodies coupled to magnetic beads as previously described (Jauregui-Lozano et al. 2021). RNA-seq, CUT&RUN, and ATAC-seq were performed in triplicate in male flies (200 flies for CUT&RUN; 400 flies for RNA-seq or combined CUT&RUN/ATAC-seq) at indicated age and ZT.

<u>RNA-seq</u>: libraries were generated from nuclear RNA using the Tecan Ovation SoLo RNA-seq library preparation kit with *D. melanogaster* AnyDeplete probes to eliminate rRNA, and sequencing was performed on the Illumina HiSeq 2000 platform.

<u>CUT&RUN and ATAC-seq</u>: bead-bound nuclei from the same biological replicate were split evenly and processed simultaneously for combined CUT&RUN and ATAC-seq to enable within-sample comparison. CUT&RUN was performed on bead-bound nuclei as described previously (Meers et al. 2019) using pAG-MNase fusion protein purified in our lab using the Pierce cobalt kit (ThermoFisher, Pierce™ His Protein Interaction Pull-Down Kit, Cat. #21227). pAG/MNase was a gift from Steven Henikoff (Addgene plasmid # 123461; http://n2t.net/addgene:123461; RRID:Addgene_123461). An IgG negative control was performed for each experimental sample (replicate) matched for antibody concentration and pAG-MNase incubation time. The following antibodies were used for CUT&RUN with the indicated pAG-MNase activation times: anti-GFP, 5 min (Abcam, ab6556); anti-H3K4me2/3, 20 min (Abcam, ab8580); anti-H3K36me3, 20 min (Abcam, ab9050); anti-H3K9me3, 20 min (Abcam, ab176916); anti-H3K27me3, 20 min (Epicypher, 13-0055); anti-H3K4me1, 30 min (Abcam, ab8895); anti-H3K4me2, 30 min (Thermo Fisher, 710796); anti-H3K4me3, 30 min (EpiCypher, 13-0041); anti-RNA Pol II-CTD , 2 min (Millipore Sigma, #050623); IgG, variable time (EpiCyper, 13-0042). The second anti-H3K4me3 (EpiCypher, 13-0041) antibody is specific for H3K4me3, whereas the H3K4me2/3 (Abcam, ab8580) although originally described as specific for H3K4me3, recognizes both H3K4me2 and H3K4me3 (Shah et al. 2018). CUT&RUN libraries were prepared using the NEBNext® Ultra™ II DNA Library Prep Kit for Illumina® with modifications to enrich for small fragments (https://dx.doi.org/10.17504/protocols.io.bagaibse) for the Clk and Cyc and Pol II CUT&RUN libraries. ATAC-seq was performed using the conditions for Omni-ATAC (Corces et al. 2017; Jauregui-Lozano et al. 2021) with recombinant Tn5 transposase from Active Motif, loaded with oligos according to the manufacturer’s protocol (Active Motif Cat. #81286). Libraries were sequenced on the Illumina NovaSeq X Plus platform. Detailed protocols for NIE-based RNA-seq, ATAC-seq, and CUT&RUN are available: dx.doi.org/10.17504/protocols.io.kxygxp1d4l8j/v3.

### RNA-seq, CUT&RUN, and ATAC-seq data analysis

We obtained at least 27 million reads for RNA-seq samples, and 7 million reads for CUT&RUN and ATAC-seq samples. All samples were normalized to library size during analysis (*i.e.,* CPM). We removed two of the RNA-seq samples (D10 ZT8 replicate 1, Set1KD ZT20 replicate 1) after preliminary analysis due to poor data quality. All other analyzed data had at least 3 biological replicates for each condition.

<u>Alignment and processing, ATAC-seq analysis</u>: Data was trimmed using Trimmomatic for paired-end reads (Bolger et al. 2014), and aligned to the *D. melanogaster* genome (BDGP6.46) using HISAT2 (Kim et al. 2019) for RNA-seq data or bowtie2 (Langmead & Salzberg 2012) for CUT&RUN and ATAC-seq data with the *--very-sensitive* (ATAC-seq, CUT&RUN) and *--dovetail* (CUT&RUN) arguments. For RNA-seq data, the first 5 bp from the forward read was removed prior to trimming using Cutadapt (Martin 2011), as recommended for Ovation SoLo libraries. SAM files were processed using SAMtools to generate sorted BAM files (Danecek et al. 2021). CPM-normalized bigWig files were generated using deepTools (Ramírez et al. 2016).

<u>RNA-seq</u>: Subread featureCounts(Liao et al. 2014) was used to count either exonic or intronic feature with intronic regions defined for the *Drosophila* genome using the *GenomicFeatures* package(Lawrence et al. 2013). We defined photoreceptor-expressed genes as having a CPM ≥ 1 in at least all 3 biological replicates. We used RUVseq(Risso et al. 2014) to account for batch effects in the knockdown RNA-seq experiments due to high number of samples. PCA plots were generated using the variance stabilizing transformation (vst function) on count data and plotPCA function in DESeq2 (Love et al. 2014). We determined relative expression values (CPM and FPKM) and identified genes that were differentially expressed across all time points (ZT) using edgeR (Chen et al. 2024; Chen et al. 2016; Robinson et al. 2009; McCarthy et al. 2012). Differential rhythmic gene expression analysis was performed using dryR (Weger et al. 2021), separately on either exonic reads or intronic reads, with categorized genes required to have BICW ≥ 0.6 for aging and BICW ≥ 0.4 for histone methyltransferase knockdown. Phase and amplitude values were also determined using dryR. All categories and BICW values are shown in Tables S1 and S3. Exon-based dryR categories were used for subsequent comparisons except for Fig. S2.

<u>CUT&RUN</u>: Peaks were called using MACS2 (Zhang et al. 2008) applying a custom-generated *Drosophila* CUT&RUN blacklist. This blacklist was created using BAM files from all IgG negative control samples from the initial Clk and Cyc CUT&RUN experiment based on a previously-described approach (Quinlan & Hall 2010). Briefly, peaks were called with MACS2 on IgG samples using a stringent FDR cutoff (0.001), and then peaks that were present in more than half of the files were selected using BEDTools (Quinlan & Hall 2010). We then merged peaks within 3000 bp of each other, and expanded these regions 1000 bp on either side to make a final BED file of blacklisted regions (Table S4). These blacklist regions were used for peak finding and subtracted for data visualization (heatmaps and metaplots). To define Clk and Cyc peak regions, we called peaks using MACS2 with an FDR of 0.05 for each replicate sample against its own IgG sample. We then selected peaks that were present in at least 2 of 3 biological replicates using the BEDTools multiinter function. Peaks within 500 bp were merged, and peaks mapped to chromosomes other than 2L, 2R, 3L, 2R, 4, X, and Y were filtered out. Blacklist regions were subtracted from these peaks to generate the final Clk:Cyc peak list. Clk:Cyc peaks were mapped to target genes based on nearest utilized TSS within 1 kb of peak (*see custom GTF generation below*).

<u>Data visualization, custom GTF, and gene ontology</u>: All CUT&RUN data is shown as CPM-normalized and subtracted for respective IgG control. Data are visualized across genomic coordinates using Gviz (Hahne & Ivanek 2016). Heatmaps and metaplots were generated using deepTools (Ramírez et al. 2016) based on a custom GTF file representing unique used transcripts. To create the custom GTF, the *Drosophila melanogaster* BDGP6.46 genome assembly release 112 was used. First, the GTF was limited to chromosomes X, 2, 3, and 4. Then, the GTF was subset to contain only unique TSSs, keeping longest transcripts. TSSs that were not used in photoreceptor cells were removed as follows: Aging RNA-seq data was counted over 5’ UTR regions using Subread featureCounts, then normalized to produce CPM using edgeR before filtering for UTR regions that had at least 1 CPM across three biological replicates. Afterward, genes that were detected using RNA-seq but did not have 5’ UTR counts (*e.g.,* noncoding RNAs) were added back into the GTF. This custom GTF is supplied as Table S4.

CUT&RUN signals at different ZTs at promoters (TSS ± 500bp) and gene bodies (500 bp downstream of TSS to TES) were calculated by averaging signal over defined region using deepTools multiBigWigSummary. Gene ontology analysis for D10 versus D50 was performed using clusterProfiler(Yu 2024; Xu et al. 2024; Yu et al. 2012; Wu et al. 2021). Gene ontology analysis for age-dependent rhythmicity categories was performed using GOrilla (Eden et al. 2009; Eden et al. 2007), and enriched GO terms presented as radial histograms showing peak expression in each genotype based on RNA-seq data using ggplot2 custom R scripts (Hadley Wickham 216AD). Custom plots were generated in R (version 4.4.1).

### Statistical analysis

Statistical analyses were performed as described in figure legends. Rhythmic gene expression analysis statistics were generated using the dryR package. Statistics for GO terms were generated using clusterProfiler. Differential gene expression statistics were generated using edgeR. Gene overlapping statistics were performed using SuperExactTest (Wang et al. 2015). Statistical significance for retinal degeneration was derived using an ANOVA with Tukey’s HSD. Statistical significance for qPCR was determined using Dunn’s test.

### Data availability

High-throughput sequencing data is available at the Gene Expression Omnibus (GEO). GSE262862: D10 and D50 RNA-seq (33 samples); GSE253177: Clk and Cyc CUT&RUN (84 samples); GSE281054: D10 and D40 H3K4me2/3, H3K36me3, H3K27me3 and H3K9me3 CUT&RUN, and D10 mCherry RNAi, Trx RNAi, Trr RNAi H3K4me1/2/3 CUT&RUN (40 samples) GSE278041: D10 and D50 ATAC-seq, RNA Pol II, H3K4me1/2/3 CUT&RUN (252 samples); GSE262534: D10 mCherry RNAi, Set1 RNAi, Trx RNAi, Trr RNAi RNA-seq (72 samples). Rhythmic expression plots for genes of interest in aging or H3K4 methyltransferase knockdown can be visualized at: https://vikkiweake.shinyapps.io/shinyr_vweake/.

### Code availability

Any custom codes used for generating the plots or analysis in this study are available upon request.

### Author contributions

S.E.M. contributed aging RNA-seq, CUT&RUN for Clk, Cyc, and RNA Pol II, and ATAC-seq. G.M. contributed HMT knockdown RNA-seq and CUT&RUN of H3K4me1, H3K4me2, and H3K4me3. M.N.M. contributed aging D10 vs D40 CUT&RUN experiments. S.A.P., S.E.M., and G.M. contributed optic neutralization experiments. S.E.M., G.M., and V.M.W. contributed the bulk of data analysis, writing, and editing.

### Competing interests

The authors declare no competing interests.

### Materials & Correspondence

Correspondence and requests for materials should be addressed to Vikki M. Weake.

## Acknowledgements

Stocks obtained from the Bloomington Drosophila Stock Center (NIH P40OD018537) were used in this study. We thank Justin Kumar, Tony Hazbun, and Paul Hardin for providing comments on this manuscript, and Daniel Mangan for assistance with Kdm5 knockdown experiments.

## Funding

Research reported in this publication was supported by the National Eye Institute of the National Institutes of Health under Award Number R01EY033734 to V.M.W. The content is solely the responsibility of the authors and does not necessarily represent the official views of the National Institutes of Health. Additional support was provided by Bird Stair funding from the Purdue University Department of Biochemistry to S.E.M. and G.M.

## Notes

### Competing Interest Statement

The authors have declared no competing interest.

