## Supplemental Figures for "Broad epigenetic shifts in the aging *Drosophila* retina contribute to its altered rhythmic transcriptome"

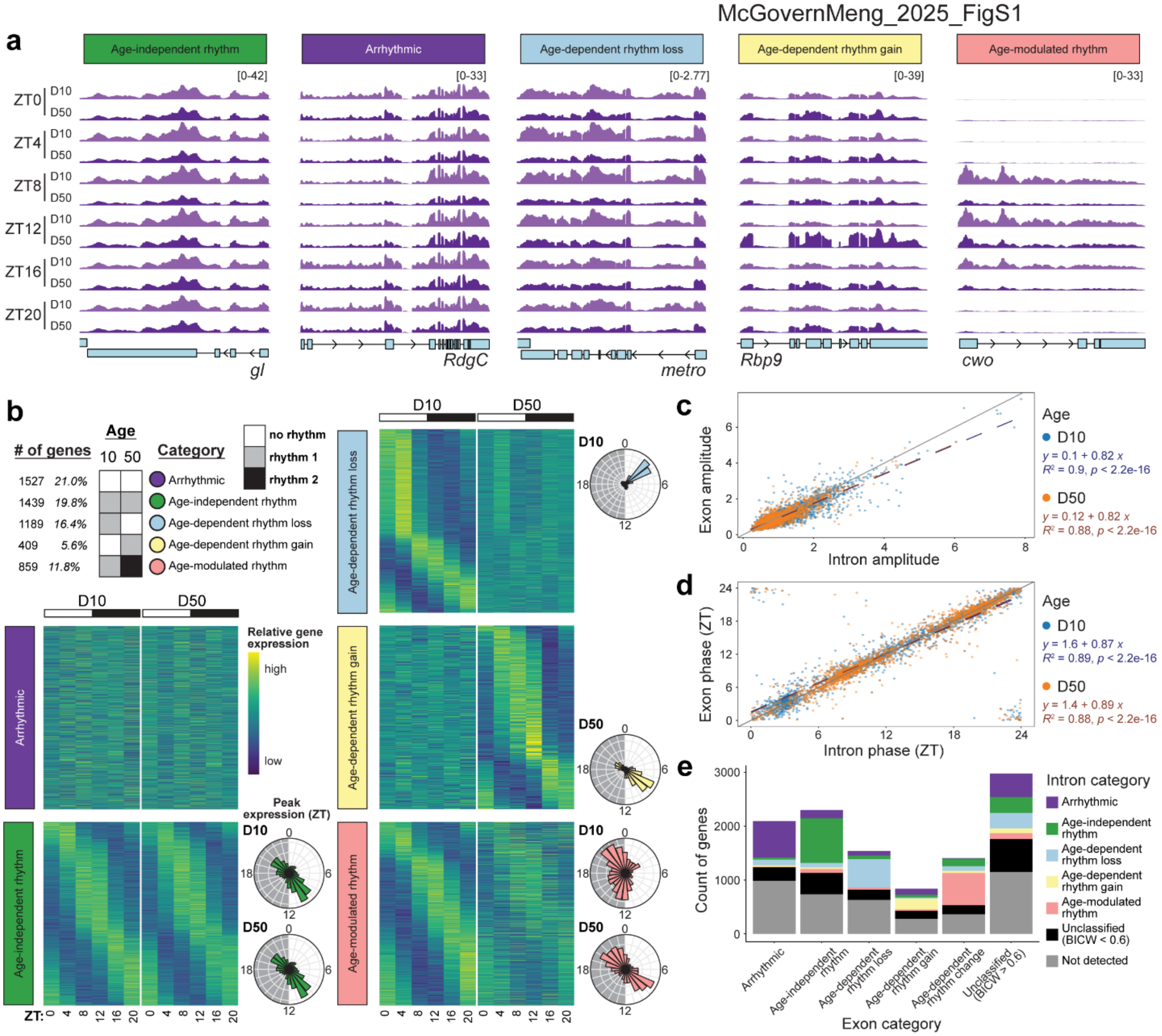


**Supplemental Figure 1: Rhythmic patterns of photoreceptor gene expression reflect nascent transcription.**

**a,** Nuclear RNA-seq data includes both intronic and exonic reads, representing nascent transcription. Tracks showing CPM-normalized RNA-seq across all ZTs at both ages for single gene examples from each aging rhythmic gene expression category. Data is mean ($n$ = 3). Exons are indicated by solid boxes and introns by connecting lines for the selected genes, labeled under each track. Arrows indicate direction of transcription. **b,** Rhythmic dryR intron-based categories (BICW ≥ 0.6). Heatmaps show $z$ score of relative expression, and radial histograms show phase of peak gene expression. **c,d,** Scatterplots comparing amplitude (**c**) or phase (**d**) using intron versus exon counts at D10 or D50. Linear regression lines (dashed) are shown for each dataset, with equations, $R^2$, and $p$ values displayed. The gray solid line represents the expected 1:1 relationship for reference. **e,** Bar plot showing the number of genes in each age-dependent rhythmicity category determined by exonic versus intronic counts. “Not detected” indicates no intronic reads were unambiguously mapped to that gene.

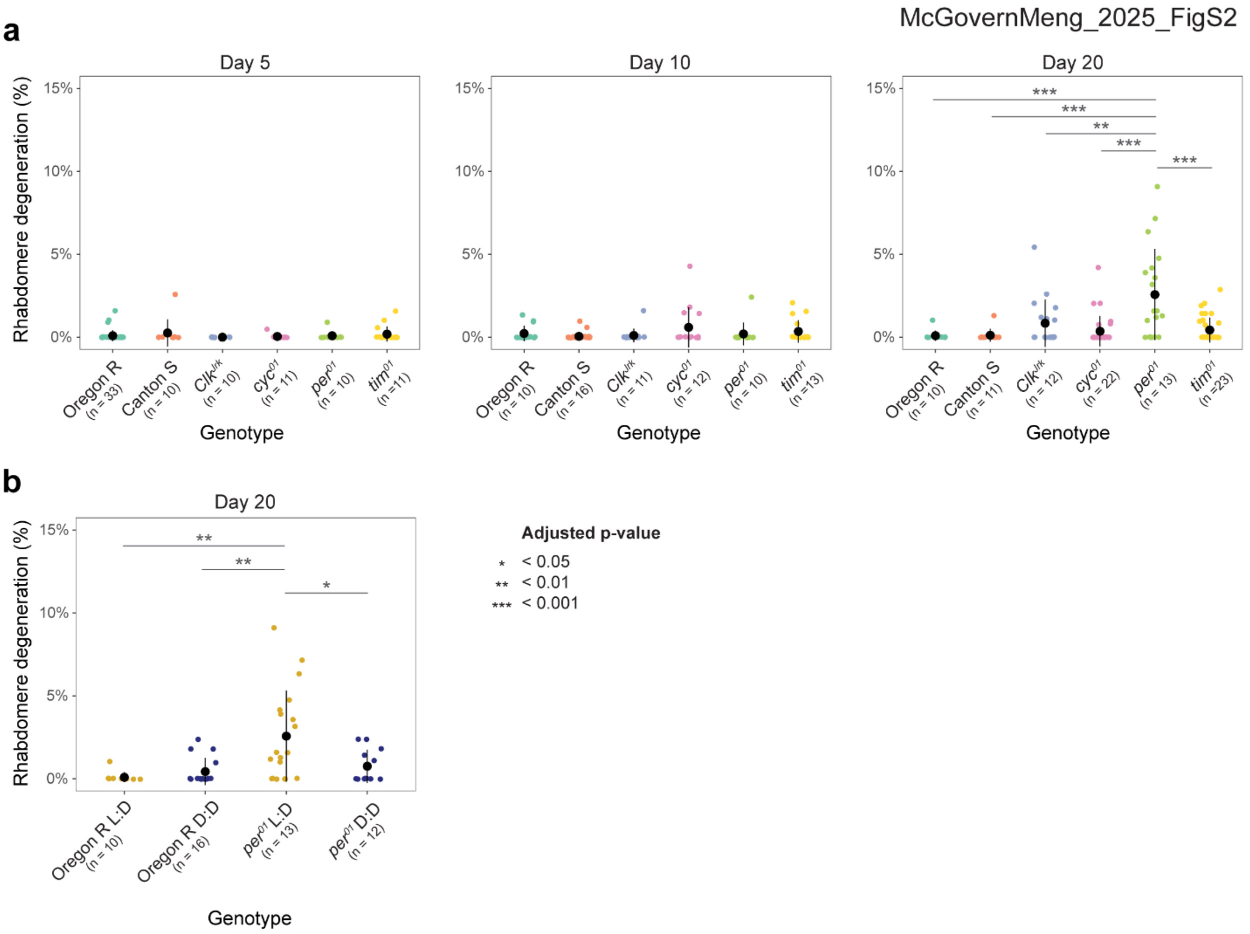


**Supplemental Figure 2: Functional *period* is required for protection against light-dependent retinal degeneration**

**a,** Retinal degeneration was assessed by optical neutralization of flies mutant for the core clock genes, *Clk*, *cyc*, *per*, and *tim* relative to the wild-type controls, Oregon R, and Canton S at Day 5, 10, and 20 of adulthood. Flies were raised in 12-hr:12-hr light:dark conditions. **b,** Optical neutralization of *per* mutant flies relative to the wild-type control, Oregon R. Flies were raised in 12-hr:12-hr light:dark (L:D) or dark:dark (D:D) conditions as indicated. * $p < 0.05$, ** $p < 0.01$, *** $p < 0.001$, ANOVA + Tukey's HSD. Number of biological replicates ($n$) is indicated per condition.

McGovernMeng_2025_FigS3

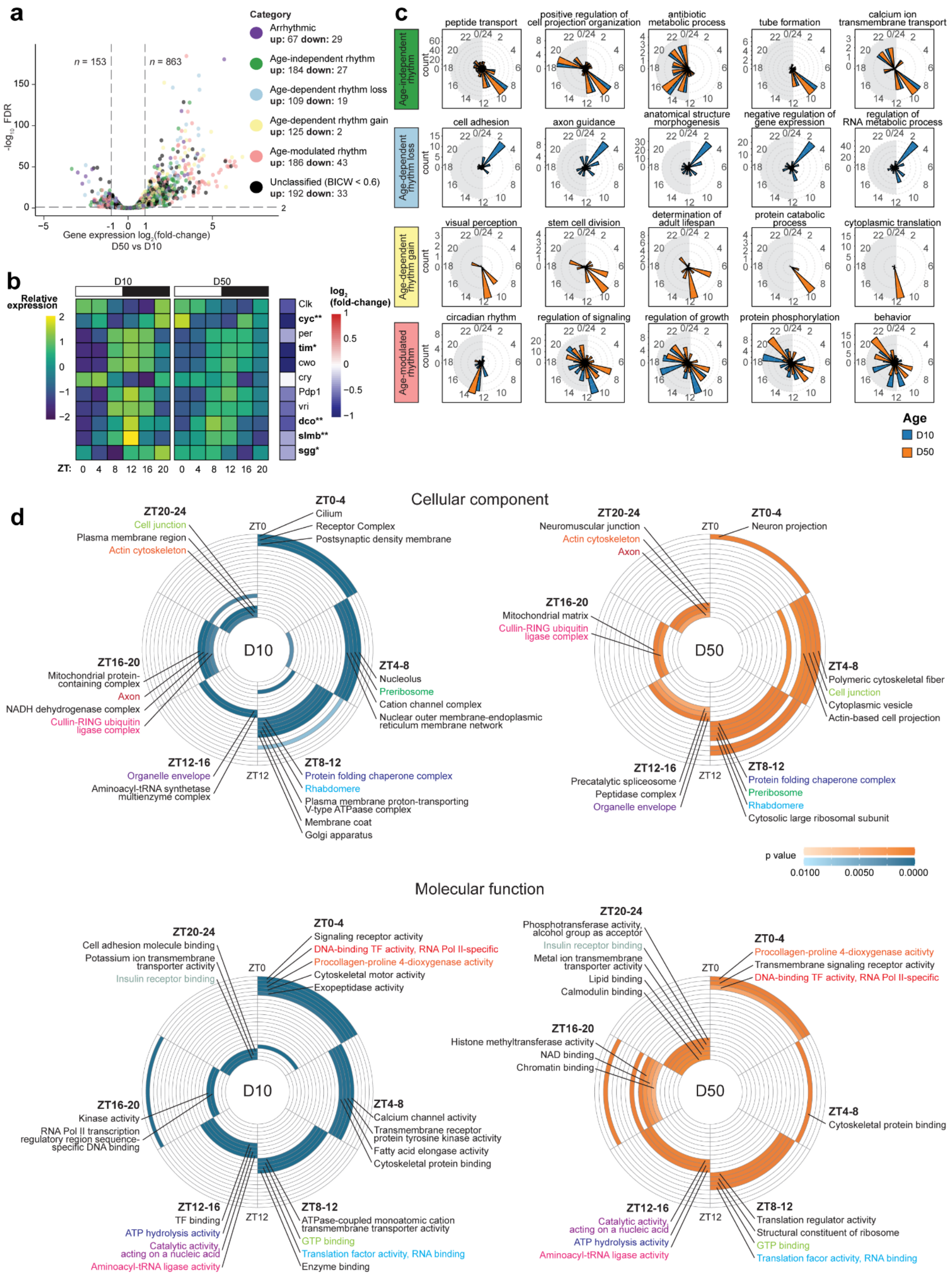

**Supplemental Figure 3: Age-dependent changes in rhythmicity do not necessarily reflect overall changes in gene expression levels.**

**a,** Differential gene expression between D10 and D50 across all ZTs was determined using edgeR, identifying 853 significantly age-upregulated genes and 153 age-downregulated genes (FDR < 0.01, $\log_2$(fold-change) ≥ 1 or ≤ -1). Individual genes are plotted as points on the volcano plot, colored by the age-dependent rhythmicity category. Numbers of overlapping genes for each comparison are shown in the legend. **b,** Heatmaps showing *z* score of relative expression for core clock genes (left) and $\log_2$(fold-change) from D10 to D50 (right). Adjusted *p* values were determined using edgeR. * $p < 0.05$, ** $p < 0.01$. **c,** Selected GO term analysis of genes in each age-dependent rhythmicity category displayed as radial histograms indicating peak gene expression phase at each age. **d,** Selected GO terms of rhythmic genes expressed most highly in four-hour intervals at D10 and D50 for cellular component (top) and molecular function (bottom). GO terms are colored by *p* value within the circles.

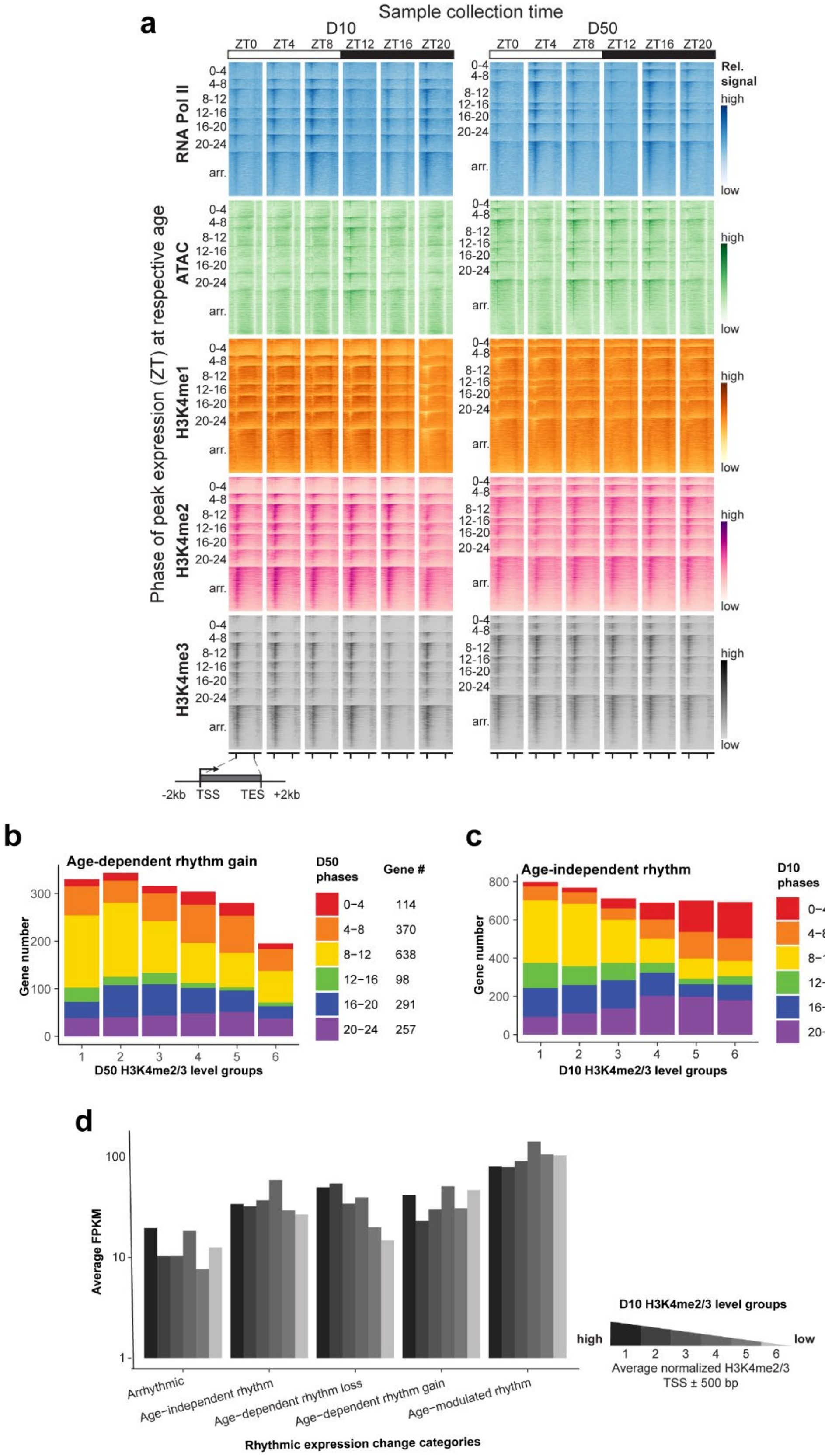
a
Sample collection time
D10
D50
ZT0 ZT4 ZT8 ZT12 ZT16 ZT20
Phase of peak expression (ZT) at respective age
RNA Pol II
ATAC
H3K4me1
H3K4me2
H3K4me3
0-4
4-8
8-12
12-16
16-20
20-24
arr.
Rel. signal
high
low
-2kb TSS TES +2kb
b
Age-dependent rhythm gain
Gene number
D50 H3K4me2/3 level groups
D50 phases
Gene #
0−4 114
4−8 370
8−12 638
12−16 98
16−20 291
20−24 257
c
Age-independent rhythm
Gene number
D10 H3K4me2/3 level groups
D10 phases
Gene #
0−4 543
4−8 551
8−12 1189
12−16 449
16−20 713
20−24 916
d
Average FPKM
Arrhythmic
Age-independent rhythm
Age-dependent rhythm loss
Age-dependent rhythm gain
Age-modulated rhythm
Rhythmic expression change categories
D10 H3K4me2/3 level groups
high
low
Average normalized H3K4me2/3
TSS ± 500 bp

40

**Supplemental Figure 4: H3K4me2/3 levels differ by both phase of peak gene expression, and age-dependent rhythmicity category**

**a,** Heatmaps representing gene bodies ± 2 kb by relative signal (*z* score) across the same gene (rows) at each ZT and age. Genes are clustered based on the phase of peak expression at D10 or D50 determined by nuclear RNA levels at each age (ZT: 0-4, 4-8, 8-12, 12-16, 16-20, 20-24, arr. = arrhythmic), and sorted within each cluster based on descending Pol II signal. **b,** Bar plot showing the phase distribution of genes that gain rhythmicity based on their relative H3K4me2 and H3K4me3 levels at D50 and expression phase at D10. Genes were separated into six H3K4me2/3 level groups ranging from highest (group 1) to lowest (group 6) based on their average normalized H3K4me2/3 scores at TSS ± 500 bp. **c,** Bar plot showing genes that maintain rhythmicity during aging, based on their phase distribution at D10. **d,** Bar plot showing average FPKM of genes in each rhythmicity category, grouped by average H3K4me2/3 scores at D10 as in **c**.

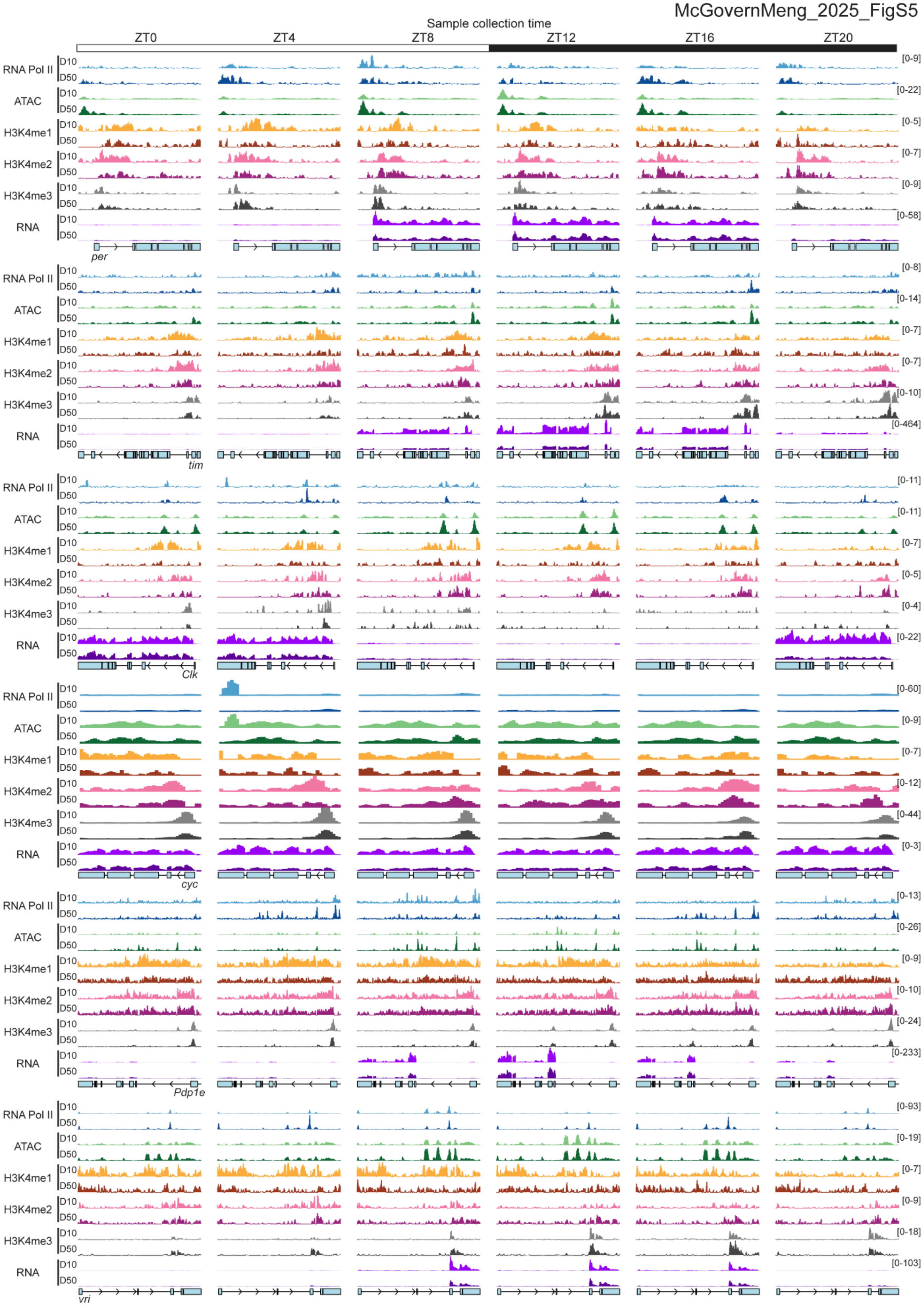
Sample collection time
ZT0
ZT4
ZT8
ZT12
ZT16
ZT20
RNA Pol II
ATAC
H3K4me1
H3K4me2
H3K4me3
RNA
D10
D50
per
tim
Clk
cyc
Pdp1e
vri
[0-9]
[0-22]
[0-5]
[0-7]
[0-9]
[0-58]
[0-8]
[0-14]
[0-7]
[0-7]
[0-10]
[0-464]
[0-11]
[0-11]
[0-7]
[0-5]
[0-4]
[0-22]
[0-60]
[0-9]
[0-7]
[0-12]
[0-44]
[0-3]
[0-13]
[0-26]
[0-9]
[0-10]
[0-24]
[0-233]
[0-93]
[0-19]
[0-7]
[0-9]
[0-18]
[0-103]

**Supplemental Figure 5: Certain core clock genes exhibit unique H3K4 methylation oscillation**

ATAC-seq, RNA-seq, and IgG-normalized CUT&RUN tracks at indicated core clock genes across ZTs at D10 versus D50. Data are mean (*n* = 3).

McGovernMeng_2025_FigS6

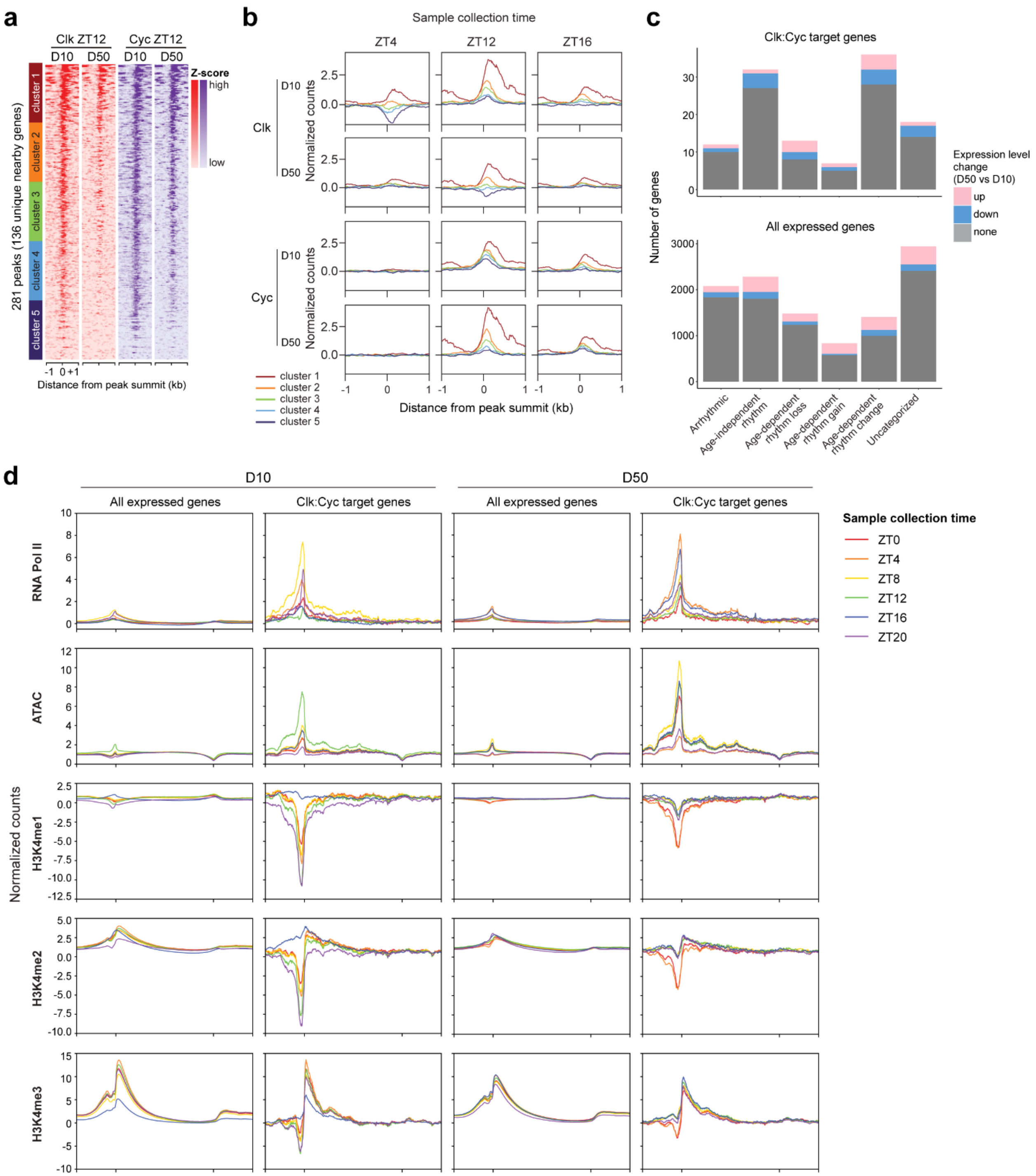


**Supplemental Figure 6: Clk:Cyc target genes have distinct epigenetic signatures**

**a,** Heatmaps representing all Clk and Cyc peak centers ± 1 kb colored by relative signal (*z* score) across the same region (rows) (*n* = 3). Heatmaps are sorted by descending Clk signal. Clk:Cyc target genes were separated into six clusters, indicated by colored bars (left), based on high (cluster 1) to low (cluster 6) Clk:Cyc signal. **b,** Gene metaplots representing mean normalized counts for Clk:Cyc peaks in each cluster. Strongest targets (cluster 1) show a distinct pattern of switching from high Clk at D10 to high Cyc at D50. **c,** Bar plot of proportion of Clk:Cyc target genes compared with all expressed genes, colored by age-dependent differential expression (≥ 1.5-fold change, adjusted $p$ value < 0.01, determined using edgeR). **d,** Gene metaplots representing mean normalized counts for CUT&RUN analysis of Pol II, H3K4me1/2/3, and ATAC-seq separated by ZT (collection time for CUT&RUN/ATAC-seq) at D10 versus D50 for all expressed genes compared to 136 Clk:Cyc target genes.

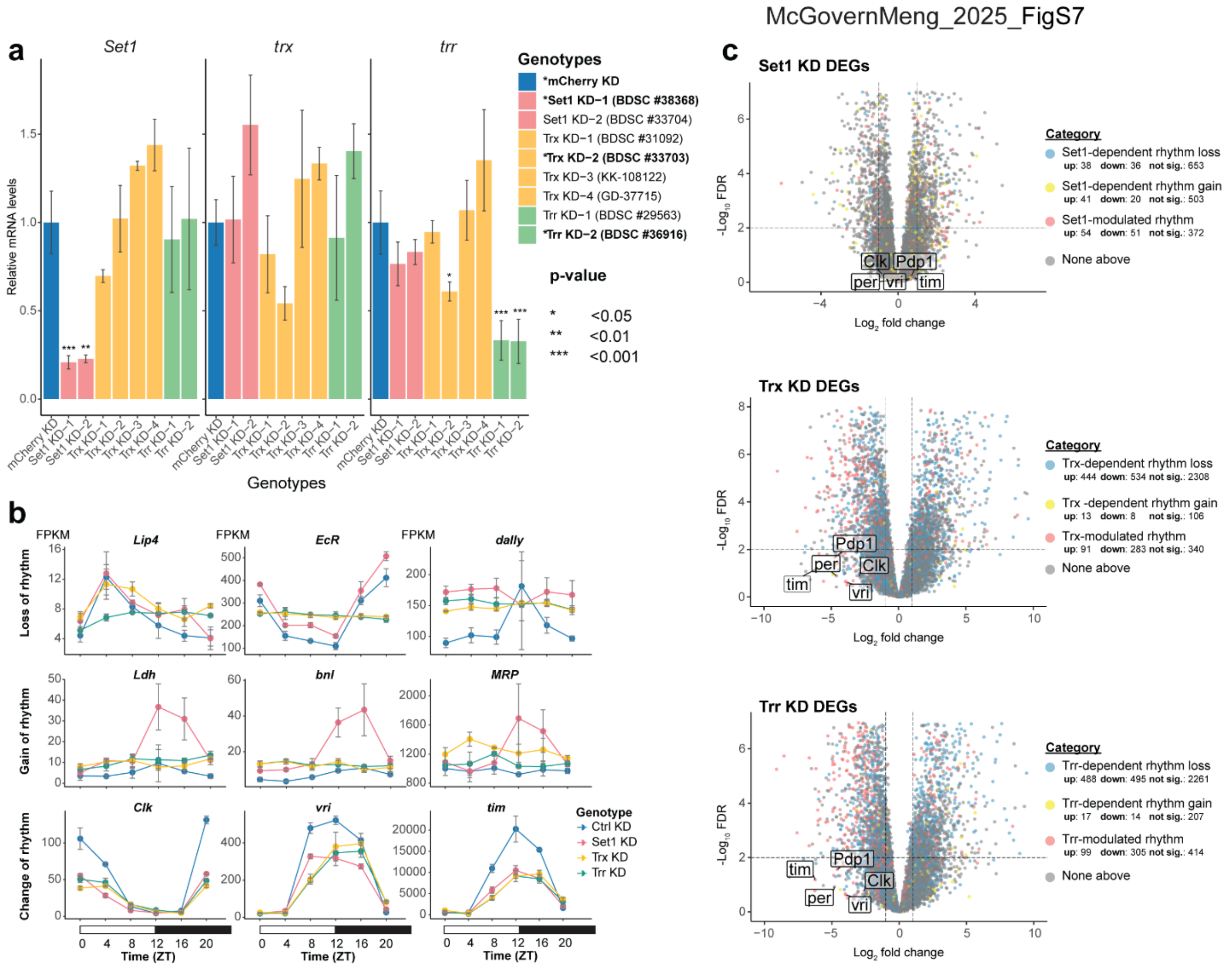


**Supplemental Figure 7: HMT-dependent changes in rhythmicity do not necessarily reflect overall changes in gene expression levels.**

**a,** qRT-PCR showing expression of *Set1*, *trr*, *trx* in indicated RNAi lines under *tub-Gal4* control. RNAi expression was restricted using $Gal80^{ts}$ by switching third instar larvae from the restrictive (18°C) to permissive (29°C) environment for 72 hours. Conditional knockdown was required because constitutive expression of some RNAi lines was lethal at the larval stage. Fly lines used in this study for further RNA-seq and/or CUT&RUN analysis are indicated in bold and marked with an asterisk. Relative mRNA levels (expression) were calculated as the geometric mean of starting quantity normalized to *Ribosomal protein L32* (*RpL32*) and *eukaryotic translation initiation factor 1A* (*eIF1A*). Bar graphs show relative expression mean ± s.d. ($n$ = 3). ** $p < 0.01$, Dunn's test. **b,** Line plots of gene expression levels for representative genes in the rhythmicity categories. Data are mean FPKM ± s.d. ($n$ = 3). **c,** Differential gene expression between Set1, Trx-, or Trr- KD vs Ctrl KD across all ZTs was determined using edgeR (FDR < 0.01, $\log_2$(fold-change) ≥ 1 or ≤ -1). Individual genes are plotted as points on the volcano plot, colored by the KD rhythmic gene expression category. Numbers of overlapping genes for each comparison are shown in the legend. Select core clock genes are labeled.

McGovernMeng_2025_FigS8

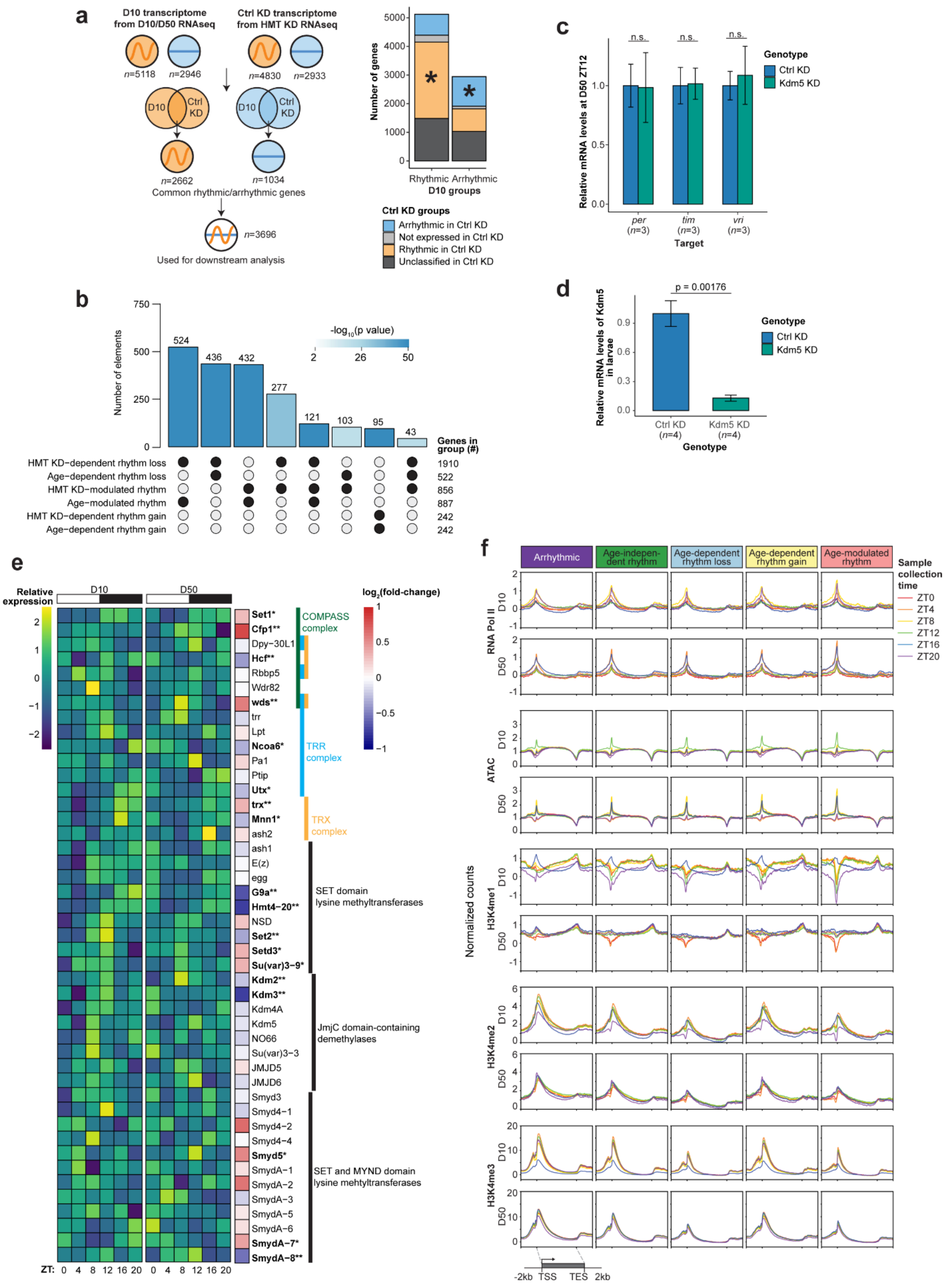

**Supplemental Figure 8: Aging and H3K4 methyltransferase knockdown comparison.** **a,** Workflow (left) to select genes for comparison between aging and HMT KD RNA-seq. Genes selected for downstream analysis were required to have rhythmicity or to be arrhythmic in both controls. Bar plots (right) show the number of genes falling into different rhythmicity categories between the two RNA-seq experiments. Asterisks mark the groups of genes that were used in the downstream analysis. **b,** UpSet plot describing the number of overlapping genes between the aging and HMT knockdown rhythmicity groups. Groups included in comparison are marked with black dots at the bottom, and bar chat on the top shows the number of overlapping genes, colored by statistical significance of intersections. **c,** qRT-PCR from adult *Drosophila* heads showing expression of *per*, *tim,* and *vri* in indicated RNAi lines under *Rh1-Gal4* control. Relative mRNA levels (expression) were calculated as the geometric mean of starting quantity normalized to *Ribosomal protein L32* (*RpL32*) and *eukaryotic translation initiation factor 1A* (*eIF1A*). Bar graphs show relative expression mean ± s.d. Sample number (*n*) is indicated. *p* values were determined using Student's t-test. **d,** qRT-PCR from *Drosophila* wandering third-instar larvae showing expression of *Kdm5* in indicated RNAi lines under *tub-Gal4* control. Bar graphs show relative expression mean ± s.d and were normalized as in **c**. Sample number (*n*) is indicated. *p* values were determined using Student's t-test. **e,** Heatmaps showing *z* score of relative gene expression (left) and $\log_2$(fold-change) from D10 to D50 (right) for selected histone methyltransferases, complex subunits, and histone demethylases. Complexes were based on FlyBase annotations. Adjusted *p* values were determined using DESeq2. * $p < 0.05$, ** $p < 0.01$. **f,** Gene metaplots representing mean normalized counts at D10 or D50 for Pol II, H3K4me1/2/3, and ATAC-seq at each of the aging rhythmic gene expression categories (columns) ($n = 3$). Lines are colored by phase of peak gene expression at the respective age.
