## Supplemental Table Legends for "Broad epigenetic shifts in the aging *Drosophila* retina contribute to its altered rhythmic transcriptome"

SIGuide.doc

**Supplemental Table 1**

D10 vs D50 photoreceptor nuclear RNA-seq processed data. DryR analysis output (DryR_D50vsD10), edgeR differential gene expression analysis (edgeR_D50vsD10), and GOrilla output for each age-dependent rhythmicity category.

**Supplemental Table 2**

Peaks of Clk:Cyc occupancy and nearest mapped genes as determined using the closest TSS that is used in photoreceptors.

**Table 3: HMT KD RNA-seq**

mCherry/Set1/Trx/Trr KD photoreceptor nuclear RNA-seq processed data. Includes DryR analysis output (DryR_HMT_KD), edgeR differential gene expression analysis (SetKD_DEGs, TrxKD_DEGs, TrrKD_DEGs).

Comparison between D10 vs D50 and mCherry/Set1/Trx/Trr KD nuclear RNA-seq processed data. Includes rhythm parameters in each condition (DryR_AgingvsHMTKD) and gene overlap analysis (AgingvsHMTKD_UpSet).

**Supplemental Table 4**

Additional methods information. Includes all genotypes used (Genotypes), all qPCR primers (Primers), the CUT&RUN blacklist generated from IgG sample data (Blacklist), and the custom GTF for unique used transcripts in photoreceptors (Custom GTF).
